# Somatic hypermutation patterns in immunoglobulin variable regions are established independently of the local transcriptional landscape

**DOI:** 10.1101/2022.05.21.492925

**Authors:** Ursula E. Schoeberl, Johanna Fitz, Kimon Froussios, Renan Valieris, Iordanis Ourailidis, Marina Makharova, Bernd Bauer, Tobias Neumann, Eva-Maria Wiedemann, Monika Steininger, Adriana Cantoran Garcia, Marialaura Mastrovito, Hugo Mouquet, Israel Tojal Da Silva, Rushad Pavri

## Abstract

Somatic hypermutation (SHM) of immunoglobulin variable (V) regions modulates antibody-antigen affinity is initiated by activation-induced cytidine deaminase (AID) on single-stranded DNA (ssDNA). Transcription is essential for SHM and AID target genes harbor activating chromatin marks and RNA polymerase II (Pol II) stalling, leading to the model that these features favor higher rates of mutagenesis. However, whether such relationships exist within V regions is undetermined. Here, we directly compared SHM and nascent transcription across four V regions and 275 non-immunoglobulin SHM targets at single-nucleotide resolution using precision run-on sequencing (PRO-seq). Although locales of Pol II enrichment and zones of Pol II stalling were detected within V regions, their correlation with SHM was not statistically significant. Moreover, SHM was robust against major reductions of activating epigenetic marks and transcription. This data suggests that SHM patterns and spectra are established independently of specific local nascent transcriptional features.

## Introduction

Somatic hypermutation (SHM) is the molecular basis for the diversification of antibodies in response to pathogens and vaccines and is hence indispensable for robust long-term immunity ^1–3^. SHM occurs in activated B cells within microanatomical structures called germinal centers in secondary lymphoid tissue upon engagement with antigens and helper T cells ^4^. Mutations are generated in the V regions of the immunoglobulin (IG) heavy (*IGH*) and light chain (*IGK*, *IGL*) genes by activation-induced cytidine deaminase (AID) ^5,6^ which acts co-transcriptionally on single-stranded DNA (ssDNA) ^7–10^. AID preferentially deaminates cytosine residues within WR<u>C</u>H motifs (where W = A or T, R = A or G and H = A, C or T) ^11^. AID also acts at the *IGH* switch regions triggering a reaction cascade leading to class switch recombination (CSR) which yields the various antibody isotypes essential for effector functions following antigen binding ^12–15^. In addition, SHM occurs at several transcribed non-IG genes and has been directly linked with B cell genome instability and cancer ^16–21^.

A longstanding question in the field is the relationship of SHM with the nascent transcriptional and epigenetic landscapes. AID is recruited to the transcriptional machinery via specific enhancers ^22–25^ and can subsequently associate with Pol II and the generic transcriptional machinery, including elongation factors ^26–30^ and RNA processing proteins ^31,32^. SHM and CSR have also been correlated with the presence of activating histone modifications typical of transcribed genes ^29,33–41^. Given that AID is a very weak enzyme ^42,43^, it has been suggested that higher rates of mutation may be favored under specific conditions that cause increased ssDNA exposure. For instance, AID can mutate both DNA strands *in vitro* when the substrate is negatively supercoiled but is far less active on relaxed substrates, leading to the idea that negative supercoils in the wake of transcribing Pol II can provide a transient source of ssDNA ^44,45^. SHM has also been correlated with features where the speed of Pol II may be reduced, resulting in longer steady-state exposure of ssDNA. These features include transcription initiation ^46^, transcription termination ^47–50^, convergent transcription ^51^, and most prominently, promoter-proximal paused Pol II and slowly elongating Pol II in gene bodies, which we refer to henceforth as Pol II stalling ^17,27,52–56^. Moreover, it has been suggested that ssDNA patches detected in V regions and non-IG SHM targets may correspond to ssDNA within the Pol II holoenzyme or from negative supercoils upstream of stalled Pol II ^54,57–59^.

The idea that stalled Pol II may play a role in SHM was first proposed nearly three decades ago ^60^. Subsequently, Pol II was shown to accumulate in IG switch regions during CSR ^52,53^ and the Pol II stalling factor, SPT5, was found to be required for efficient AID targeting to IG switch regions and non-IG AID target genes ^27^. Pol II accumulation in V regions was also found to be higher in SHM-competent germinal center B cells but not in SHM-incompetent primary B cells ^54^. Recently, studies using model reporter substrates have found that SHM driven by mutation enhancers correlated with an increase in the frequency of ssDNA patches along with increased SPT5 and Pol II stalling ^55^.

Indeed, high levels of Pol II and SPT5 were found to be sufficient to identify new AID target genes via a machine learning approach ^17^. Collectively, this leads to a model suggesting that the likelihood of SHM is higher in the vicinity of zones of stalling and/or convergent transcription due to increased time and access to ssDNA, relative to the neighboring DNA lacking any such feature ^51,55,61–63^.

Importantly, however, the nascent transcriptional landscape of V regions has not been determined and hence a systematic comparison between transcriptional and SHM patterns is lacking. There is evidence for Pol II stalling in V regions based on the accumulation of Pol II observed with chromatin immunoprecipitation followed by quantitative PCR (ChIP-qPCR) ^48,54^. However, due to the very short length of V regions (∼500 bp) coupled with the low resolution of ChIP-qPCR and ChIP-sequencing (ChIP-seq), one cannot ascertain if Pol II is stalled across the entire V region or at specific locations, and therefore, whether sites of recurrent stalling correlate with zones of increased mutagenesis.

Here, we address this question by using precision run-on sequencing (PRO-seq) and the related method, PRO-cap, in combination with a customized bioinformatics pipeline to map reads to V regions. By identifying the precise 3’ end of nascent RNA within engaged Pol II complexes, PRO-seq provides a map of the location and orientation of actively transcribing Pol II genome-wide at single nucleotide resolution ^64,65^. PRO-cap identifies the 5’ ends of capped nascent RNAs that are associated with engaged Pol II molecules and hence provides maps of the location and orientation of transcription initiation sites at single nucleotide resolution ^64,66^. For these reasons, PRO-seq and PRO-cap are amongst the most widely used assays for studying nascent transcriptional profiles genome-wide and they have been particularly important in identifying recurrent sites of Pol II stalling and initiation at promoters, enhancers and within gene bodies ^66–72^. The single nucleotide resolution of the assay means that PRO-seq can identify recurrent sites of Pol II stalling within the short V regions. With this approach, we could ask whether and to what extent the discrete SHM patterns in V regions and non-IG SHM targets can be plausibly explained by recurrent stalling, initiation or convergent/antisense transcription. The same approach was used to study the relationship between the levels of activating chromatin marks with SHM within V regions. We found weak and inconsistent correlations between mutation rate and any of these co-transcriptional features or epigenetic marks at all four human and mouse V regions analyzed as well as 275 previously-identified AID target loci in mice ^17^. Thus, despite the global correlation between AID targeting and such features, when probed at the single nucleotide level, there does not appear to be a strong relationship between them, raising the possibility that ssDNA associated with these transcriptional features may not be the major source of ssDNA for AID.

## Results

### V gene locales harbor an atypical chromatin landscape

As a model system of SHM, we chose Ramos, an IgM-positive human Burkitt lymphoma-derived cell line where the single functional, pre-recombined *IGH* V region consists of the V gene, VH4-34, the diversity segment, DH3-10, and the joining segment, JH6. Ramos cells constitutively express AID and undergo low levels of SHM mostly at C:G residues ^73,74^. However, SHM can be significantly boosted in these cells upon overexpression of AID variants, and widely used for the study of SHM^25,26,33,34,39,48,55,57,58,73–78^.

To determine the chromatin landscape of the Ramos *IGH* locus, we performed chromatin immunoprecipitation followed by quantitative PCR (ChIP-qPCR) for histone modifications associated with active promoters and enhancers (histone H3 acetylated on lysine 27, H3K27ac or trimethylated on lysine 3, H3K4me3) and transcription elongation in gene bodies (H3 trimethylated on lysine 36, H3K36me3). At active genes, H3K27ac and H3K4me3 are enriched at nucleosomes flanking the transcription start site. Surprisingly, given that *IGH* is highly transcribed, we observed that the V region is characterized by relatively low levels of all marks that, importantly, was not due to decreased nucleosome occupancy as judged by the levels of histone H3 (Supplementary Fig. S1A). As a result, the first nucleosome from the transcription start site has the lowest levels of H3K27ac and H3K4me3 which is contrary to what is usually observed at highly active genes where the first nucleosome harbors the highest levels of these marks. In contrast, much higher levels of all marks were observed in the intronic regions flanking the Eμ enhancer with an expected decrease at Eμ itself where nucleosomes are occluded by the presence of transcription factors (Fig. S1A) ^79^. Importantly, these atypical profiles were also observed at the murine *Igh* in primary B cells expressing the B1-8^hi^ V region and activated with lipopolysaccharide (LPS), interleukin 4(IL4) and RP105 ^80^ (Fig. S1B). We conclude that V regions are marked by relatively low levels of activating histone marks and that this landscape is conserved between the human Ramos and mouse B1-8^hi^ V regions.

### The Eμ enhancer regulates the levels of chromatin marks in the flanking intronic DNA but not within the V region

The fact that the highest enrichments of chromatin marks were seen on either side of the Eμ enhancer suggested that this element may be responsible for the deposition of these marks. This raised the question of whether deletion of Eμ (Eμ^−/−^) would alter the levels of these marks and affect SHM. SHM results in the loss of WR<u>C</u>H motifs (hotspot saturation) as well as a gain of new hotspot motifs ^81^, hence continuous SHM in AID-sufficient Ramos cells can confound subsequent analysis of SHM profiles. To ensure that all cells have the same V gene sequence prior to SHM, we ablated *AICDA*, encoding AID, in Ramos cells (AID^−/−^) and subsequently deleted Eμ to obtain AID^−/−^ Eμ^−/−^ cell lines (Fig. S1C). Specifically, we used CRISPR-mediated editing to delete a 583 bp region corresponding to the peak of chromatin accessibility measured by assay for transposase-accessible chromatin (ATAC-seq) at Eμ in AID^−/−^ cells (AID^−/−^ Eμ^−/−^) (Fig. S1C). Based on PRO-cap analysis of transcription start sites (see below), this deletion eliminates the Iμ promoter that encodes the sterile germline *IGHM* transcript and antisense transcription start sites residing in Eμ (Fig. S1C). We chose two independent clones, AID^−/−^ Eμ^−/−^ c1 and AID^−/−^ Eμ^−/−^ c2, for subsequent experiments (Fig. S1D).

To comprehensively visualize profiles of these histone marks across the V region and flanking sequences, we performed ChIP followed by deep sequencing (ChIP-seq) for H3K27ac, H3K4me3 and H3K36me3. We generated a custom chromosome spanning the Ramos V region and promoter. Importantly, the Ramos V region is highly mappable and hence there is minimal loss of reads due to multimapping with other V gene families (see later section). The results in AID^−/−^ Ramos cells were consistent with ChIP-qPCR in that the levels of all marks were lowest in the promoter-proximal region and gradually increased into the V region, peaking in the intronic sequences flanking Eμ (Fig. 1A and B). This analysis also revealed the absence of bimodal distributions of H3K27ac and H3K4me3 at the VH4-34 promoter, which is commonly observed at highly active divergently transcribed genes. In AID^−/−^ Eμ^−/−^ cells, all three marks were decreased (Fig. 1 A and B), a finding that we confirmed by ChIP-qPCR, and which was not due to changes in nucleosome occupancy as determined by histone H3 measurements (Fig. 1C). Using antibodies against pan-H3 and pan-H4 acetylation, we observed a significant decrease in histone acetylation across the locus in AID^−/−^ Eμ^−/−^ cells (Fig. 1C). Moreover, another mark of transcription elongation, H3K79me3, was also reduced in AID^−/−^ Eμ^−/−^ cells (Fig. 1C). All marks in AID^−/−^ Eμ^−/−^ cells were significantly reduced up to 4-fold in the intron where they are normally at their peak levels (Fig. 1 A to C). However, within the V region, where these marks are normally at their lowest levels, there was no significant change in AID^−/−^ Eμ^−/−^ cells Fig. 1, A to C).

**Figure 1:**
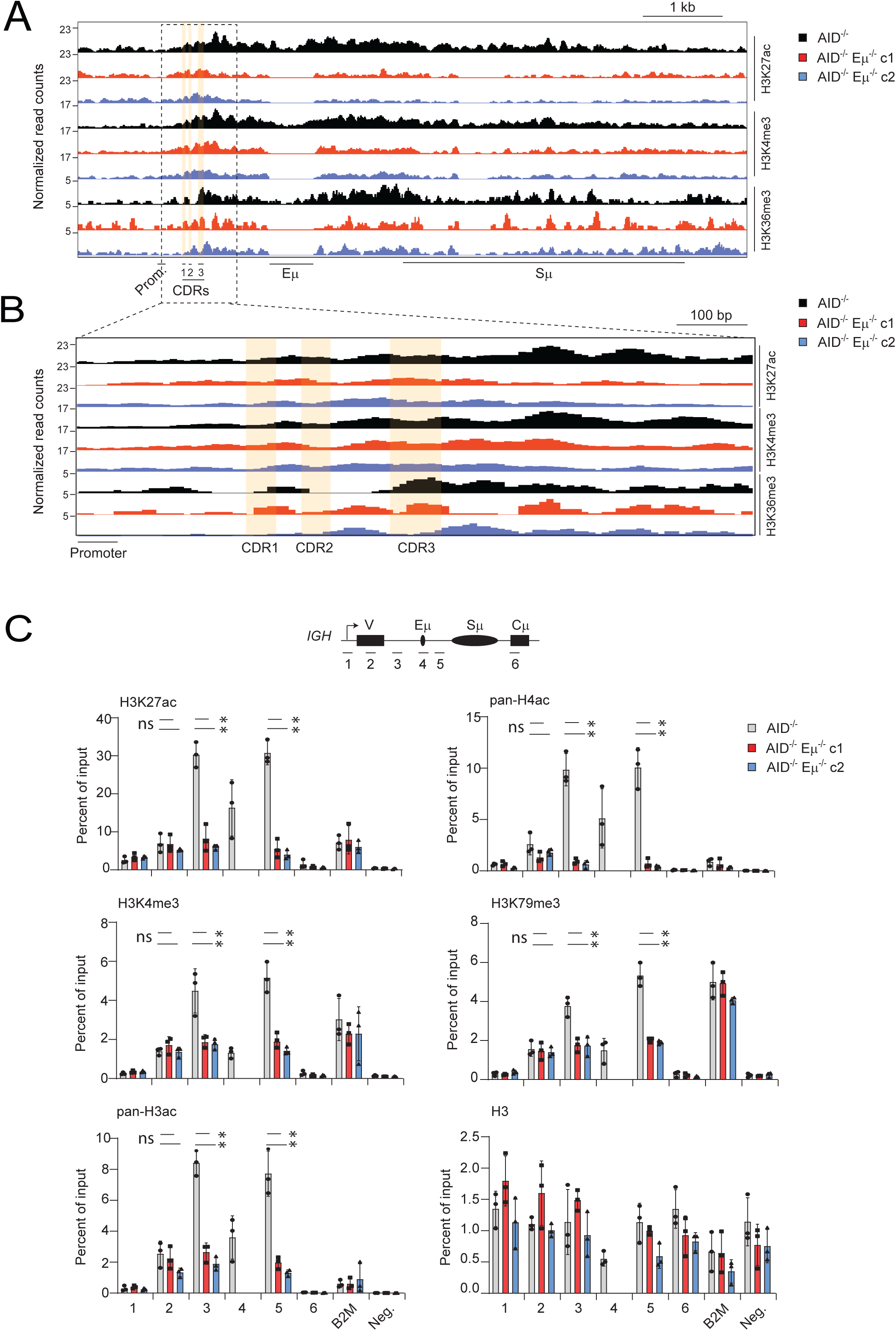
The Eμ enhancer regulates the chromatin landscape in the intron but not in the V region in Ramos human B cells. **(A)** ChIP-seq profiles of H3K27ac, H3K4me3 and H3K36me3 at the *IGH* locus from AID^−/−^ and two independent clones (c1 and c2) of AID^−/−^ Eμ^−/−^ Ramos cells, the latter generated as described in Fig. S1C-D. The sequence annotations indicate the positions of the *IGH* promoter (Prom), the complementary determining regions (CDR1-3), the intronic enhancer (Eμ) and the switch μ region (Sμ). The bioinformatic approach to include multimapping reads (explained in the text and depicted in Fig. 3A) was applied. **(B)** A magnified view of the locus from A. **(C)** ChIP-qPCR analysis at *IGH* from AID^−/−^ and AID^−/−^ Eμ^−/−^ Ramos cells (c1, c2) to measure the relative levels of H3K27ac, H3K4me3, H3K79me3, pan-acetylation of histones H3 and H4, and histone H3. The amplicons used for PCR are indicated in the schematic diagram at the top. The active *B2M* gene is used as a positive control and a gene desert segment on chromosome 1 is used as a negative control (Neg.). The data represent three independent experiments. Asterisks indicate *P* < 0.05 using the Student’s t-test and ns indicates not significant (*P* > 0.05).

Consequently, AID^−/−^ Eμ^−/−^ cells harbor a chromatin landscape wherein the levels of all marks appear to be comparable between the V region and the intron (Fig. 1C, compare the measurements from amplicons 2, 3 and 5 in AID^−/−^ and AID^−/−^ Eμ^−/−^ cells). We conclude that Eμ is responsible for the higher levels of chromatin marks within the intron but does not influence the deposition of these marks within the V region.

These changes in the chromatin landscape in AID^−/−^ Eμ^−/−^ cells were accompanied by a 2-4-fold decrease of nascent transcription in the V region and intron, as well as a 2-fold decrease of the spliced *Igh* mRNA (Fig. 2 A and B). The decrease in transcription within the V region and intron implies that Eμ regulates optimal transcription initiation from the V gene promoter. Nevertheless, despite these decreases in *Igh* transcripts, surface IgM levels were unaffected in Eμ^−/−^ cells suggesting a lack of absolute correlation between mRNA and protein levels (Fig. S1E). We conclude that Eμ positively and significantly regulates nascent transcription in the V region but has no significant impact on its epigenetic landscape.

**Figure 2:**
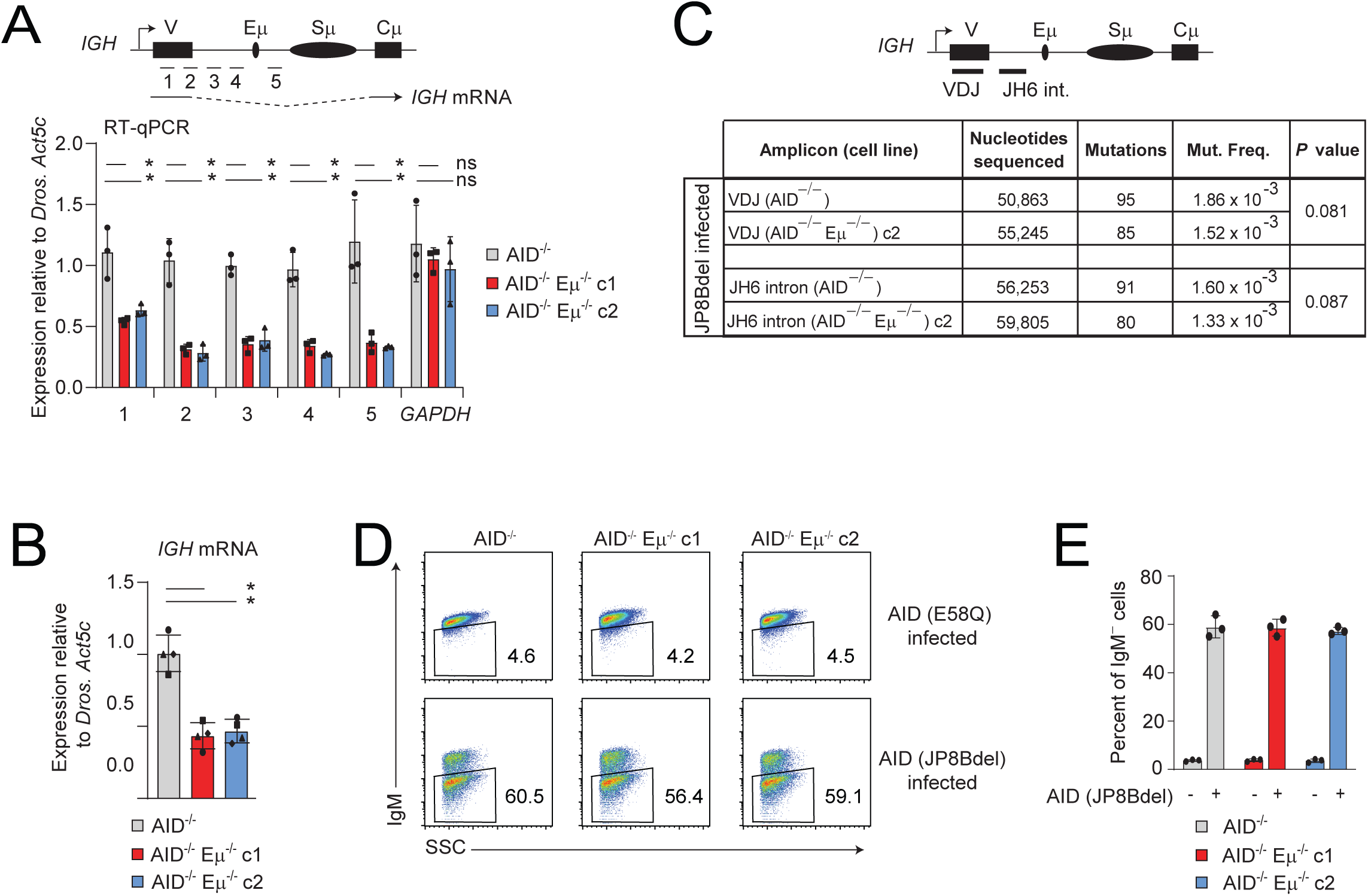
Ablation of the Eμ enhancer significantly decreases nascent transcription but not SHM. **(A)** RT-qPCR measurements of nascent transcripts at the Ramos V region in AID^−/−^ and AID^−/−^ Eμ^−/−^ cells (c1, c2). To account for potential clonal variation, Ramos cells were spiked with *Drosophila* S2 cells prior to RNA extraction. The data was normalized to the levels of the *Drosophila* housekeeping gene, *Act5c*. *GAPDH* mRNA was used as a control. Asterisks indicate *P*<0.05 using the Student’s t-test and ns indicates not significant (*P* >0.05). **(B)** RT-qPCR analysis as in A measuring the spliced *IGH* mRNA. **(C)** Table of mutation frequencies at the Ramos V region and the JH6 intron (amplicons shown in the diagram above the table) in AID^−/−^ and AID^−/−^ Eμ^−/−^ cells following infection with AID(JP8Bdel) for 7 days. Statistical analysis was performed with the Student’s t-test. **(D)** Flow cytometry analysis of IgM expression in AID^−/−^ and AID^−/−^ Eμ^−/−^ Ramos cells infected with AID(JP8Bdel) for 7 days. **(E)** Bar graph summarizing the flow cytometry analyses from D, showing the percent of IgM loss from three independent experiments.

### SHM frequency does not correlate with the levels of nascent transcription or activating histone marks in the V region locale

AID^−/−^ Eμ^−/−^ cells provide an ideal system to ask whether changes in the levels of nascent transcription and chromatin marks directly impact on SHM without any contribution from secondary, indirect effects. Hence, we sequenced the V region as well as the JH6 intron immediately downstream (Fig. 2C, upper panel). This analysis allowed us to compare SHM between a region where chromatin marks were normally low and Eμ-independent (V region) and a region where the marks were normally high and strongly Eμ-dependent (JH6 intron). To measure SHM, we infected AID^−/−^ and AID^−/−^ Eμ^−/−^ cells with AID (JP8Bdel), a C-terminal truncation mutant of AID that results in nuclear retention and major increase in mutation rates after 6 days ^82^. As a control, we also infected cells with a catalytically inactive AID mutant (E58Q) ^82^. As discussed in a later section, similar SHM spectra and profiles were obtained upon infection with a hyperactive mutant of AID (m7.3), which retains the C terminal domain ^83^. However, since AID (m7.3) required much longer culturing post-infection (21 days), we used AID (JP8Bdel) for our analyses.

In both the V region and JH6 intron, we observed a small (∼20%) decrease in SHM frequency in AID^−/−^ Eμ^−/−^ relative to AID^−/−^ cells expressing AID (JP8Bdel) that was not statistically significant (Fig. 2C). This result is consistent with mouse studies where loss of Eμ caused only modest decreases in SHM ^84,85^. As an alternative readout of SHM activity, we assayed for the loss of surface IgM, which occurs due to AID-induced stop codons or frameshifts ^73^. The results showed that the magnitude of IgM loss was similar between the control and Eμ^−/−^ cells suggesting that the minor decreases in SHM do not translate into equivalent changes in surface IgM (Fig. 2 D and E). We conclude that the Eμ enhancer is dispensable for robust SHM in Ramos cells.

By comparing the levels of chromatin marks with SHM frequencies in the V region and JH6 intron, we can draw three major conclusions: First, we infer that the levels of SHM do not correlate with those of activating histone marks. Indeed, SHM frequency is slightly lower in the intron (1.6 x 10^-3^), which harbors the highest level of these marks, than in the V region (1.86 x 10^-3^), which has the lowest levels of these modifications (Fig. 2C). Second, as exemplified by the JH6 intron, SHM is resistant to major decreases in these marks. Third, we attribute the small reduction in SHM in AID^−/−^ Eμ^−/−^ cells to the decrease in nascent transcription although, here too, the decrease of SHM (∼20%; Fig. 2C) is considerably smaller than that of nascent transcription which is reduced up to 70% in the JH6 intron of AID^−/−^ Eμ^−/−^ cells (Fig. 2A, amplicons 2-5).

We conclude that although epigenetic modifications mark loci undergoing SHM and may contribute to SHM, their levels do not correlate with SHM frequencies and their significant reduction does not have a major impact on mutation rates, at least in this system.

### Elucidating the nascent transcriptional landscape of the Ramos V region

We next asked whether SHM patterns are dictated by local features of nascent transcription landscape that could transiently lead to increased ssDNA exposure, such as during transcription initiation, stalling and/or convergent/antisense transcription. Since V genes occur as families with varying degrees of sequence identity, mapping of next-generation sequencing (NGS) reads to recombined V regions using standard alignment workflows eliminates reads which map to more than one location in the genome (multimappers) and only align reads mapping uniquely (typically allowing 2-3 mismatches). In Ramos cells, visual inspection of PRO-seq tracks on the UCSC genome browser revealed robust coverage within the VH4-34 gene with uniquely-mapping reads indicating that this V gene is highly mappable (not shown). To address this more systematically, we retrieved the normally discarded, multimapping reads and allowed them to re-align to the recombined Ramos V region with the important proviso that they only multimap to annotated V, D or J segments and not elsewhere in the genome (Fig. 3A). Separate tracks were created for uniquely mapping reads and the recalled multimapping reads, which were then combined to obtain the total profile (Fig. 3B).

**Figure 3:**
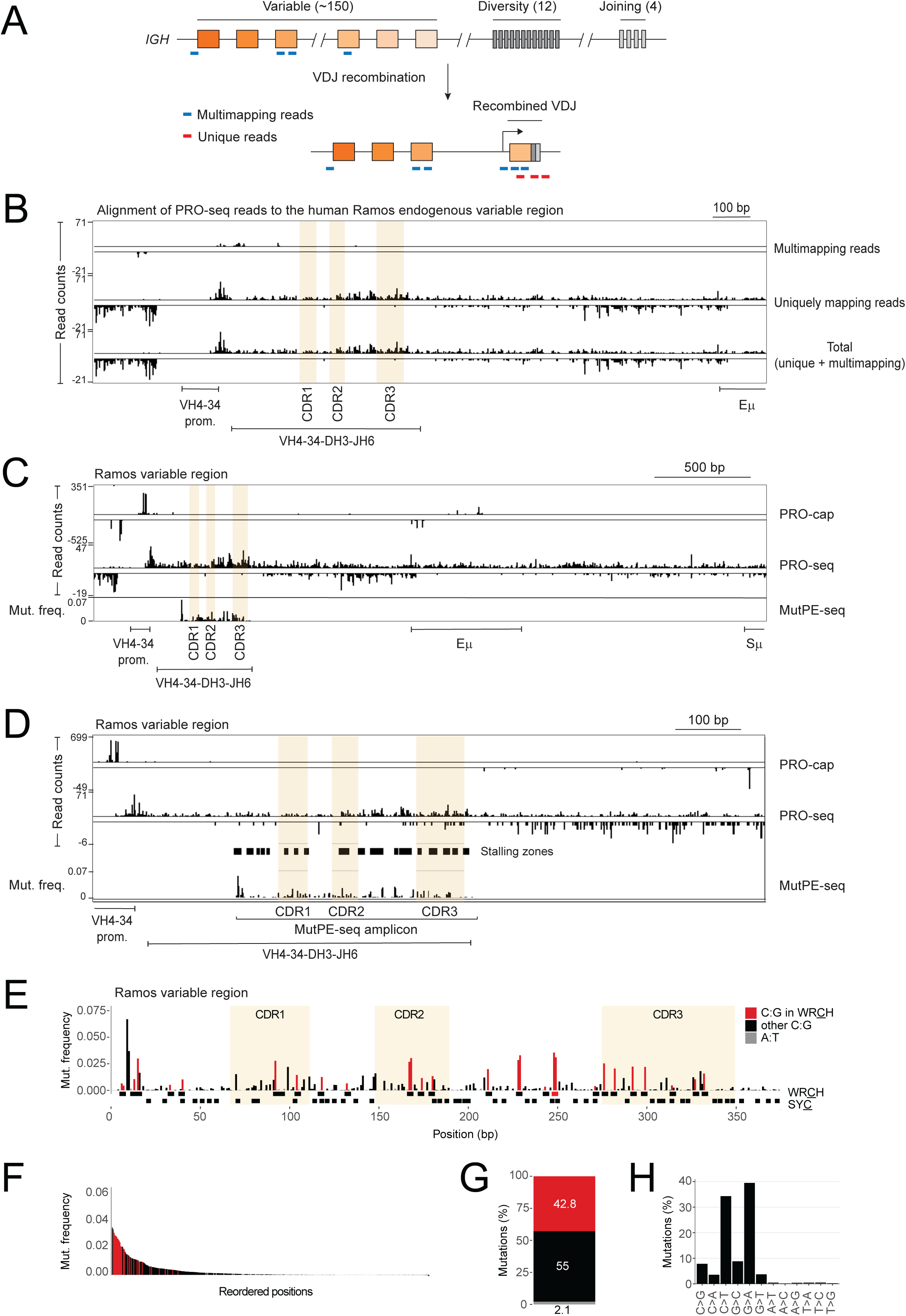
Comparison of transcriptional and mutational landscapes at the endogenous Ramos V region. **(A)** Scheme depicting the strategy used to align multimapping reads to the V region. As described in the text and Methods, since upstream, non-recombined V genes are silent in Ramos cells, a read that maps to the recombined V region is retained if it is mapping to any V, D or J segment of the human *IGH* locus but nowhere else in the human genome. The same principle is applied when mapping reads to murine V regions to the mouse genome. **(B)** Integrative Genomics Viewer (IGV) browser snapshot of nascent RNA 3’ ends (by PRO-seq) aligned to the Ramos V region. Multimapping (top track) reads are separated from uniquely mapping reads (middle track), and these two tracks are subsequently combined to generate the total profile (bottom track. **(C)** PRO-cap and PRO-seq 5’ and 3’ end densities, respectively, at the *IGH* locus along with mutation frequencies at the V region displayed on the Integrative Genomics Viewer (IGV) browser. Mapping of the V region transcriptome was done via the bioinformatic pipeline outlined in A and exemplified in B (total profiles are shown). The locations of the antigen-binding complementary determining regions (CDR1-3) are highlighted. Mutation analysis via MutPE-seq was performed following infection of AID^−/−^ cells with AID(JP8Bdel)-expressing lentiviruses for 7 days. **(D)** A magnified view of the V region from C above. The stalling zones are shown in the panel between the PRO-seq and MutPE-seq tracks. **(E)** Details of the mutation patterns at the Ramos V region. Mutated cytidines in AID hotspot motifs (WR<u>C</u>H) are displayed as red bars. All other C:G mutation are shown as black bars and A:T mutations as grey bars. The panel under the graph shows the position of both hotspot (WR<u>C</u>H in black, AGCT in red, upper panel) and coldspot (SY<u>C</u>, bottom panel) motifs. **(F)** Waterfall plot with mutations ordered from highest (left) to lowest (right) frequency following the color code described in E. **(G)** Mutation frequency bar plot showing the percentages (indicated within the bars) of the three mutations classes following the color code described in E. **(H)** Bar graph indicating the percentage of the type of mutation indicated on the X axis. The C to T and G to A transition mutations are the signature of AID activity.

We measured nascent transcriptional activity with PRO-seq and capped initiation sites with PRO-cap^66^. For simplicity, we refer to relative accumulation of PRO-seq signal within the gene body as Pol II stalling or stalled Pol II in order to differentiate it from promoter-proximal pausing which is also reported by PRO-seq as the enrichment of signal typically 40-80 bp from transcription start sites. Most notably, promoter-proximal pausing occurs via the binding of NELF, which inhibits elongation whereas elongating Pol II lacks NELF and contains additional proteins such as SPT6 and the PAF complex ^86–89^. Accumulation of Pol II signal in gene bodies may arise due to the slowing down of Pol II as a result of steric barriers such as secondary structures or premature termination signals. However, initiation events within gene bodies could also give rise to a paused Pol II state akin to promoter-proximal pausing and can be identified if PRO-seq accumulation is associated with upstream PRO-cap enrichments.

Our mapping pipeline revealed that the Ramos V region is predominantly covered by uniquely-mapping reads with very few multimapping reads (Fig. 3B). Of note, the same pipeline was used for the ChIP-seq analysis of Ramos cells described in Fig. 1 above. Therefore, this approach allowed us to delineate the transcriptional and epigenetic landscapes of IG V regions. In contrast to the lack of a bimodal peak of activating histone marks (Fig. 1A), the Ramos VH4-34 promoter showed clear bidirectional transcription and promoter-proximal pausing, the latter being noticeable as the PRO-seq enrichments shortly downstream of the promoter-associated PRO-cap initiation signals (Fig. 3C). Next, we infected cells with AID (JP8Bdel) and analyzed SHM patterns in the V region using an optimized version of mutation analysis with paired-end deep sequencing (MutPE-seq) of a PCR-amplified fragment corresponding to the V region ^90^ (Fig. 3E). Mutations occurred robustly at C:G residues within WR<u>C</u>H motifs with weak targeting of A:T residues, as expected from Ramos cells. Moreover, the cold spot motif, SY<u>C</u> (where S = C or G and Y = C or T) was poorly mutated (Fig. 3 F-H). A similar mutation profile and spectrum was observed when cells were infected with the hyperactive AID mutant, AID (m7.3) ^83^, for 21 days, indicating that the absence of the C terminal sequence in AID (JP8Bdel) does not alter its targeting to the V region or its motif preferences (Fig. S2A-D).

Visual comparison of the profiles of SHM and nascent transcription at the Ramos V region showed that some residues with relatively high mutation rates were associated with increased PRO-seq densities in the neighborhood, such as in the framework region between CDR2 and CDR3 (Fig. 3D). However, other mutations did not occur in the vicinity of higher PRO-seq signals or internal initiation events as measured by PRO-cap. For example, the region of highest PRO-seq read density at the 3’ end of the V region is not preferentially mutated over upstream C:G motifs where PRO-seq signal is much weaker (Fig. 3 D and E). Indeed, the most mutated residue is located near the 5’ end of the V region where nascent transcription signals are lower than the 3’ end (Fig. 3D).

### SHM frequency does not significantly correlate with Pol II stalling or transcriptional strength at the Ramos V region

To statistically evaluate the relationship between mutations and Pol II stalling, we mapped stalling sites as per the definition used in previous studies, namely, as positions covered by at least five PRO-seq reads and where the signal is >3 standard deviations greater than the mean of the flanking 200 bp region ^91^ (see Methods). Stalling sites were extended by 3 bp on either side to create 7 bp stalling zones (Fig. S3A). This window size was chosen to reflect the approximate length (6-8 bp) of the transcription bubble in elongating Pol II complexes ^88,89^. Next, overlapping stalling zones were merged and only those stalling zones lying within the MutPE-seq amplicon (displayed as black boxes in Fig. 3D) were considered for further analysis. All regions within the MutPE-seq amplicon that were not defined as stalling zones were termed non-stalling zones. Finally, the normalized mutation rate (mutations/width of zone) was calculated for each stalling and non-stalling zone (Fig. S3A). Mutation frequencies between stalling and non-stalling zones were compared, and statistical significance determined by the Wilcoxon rank-sum test (Fig. S3B). The results showed no statistically significant difference in mutation frequency between stalling and non-stalling zones (*P* = 0.56, Fig. S3B). Importantly, similar results were obtained when the analysis was performed with larger stalling zones (up to 21 bp).

As an alternative to identifying stalling zones, we asked whether C:G residues with higher mutation frequencies occurred in locales having higher PRO-seq densities, that is, whether mutation frequency corelated with local transcriptional strength. We divided all C:G residues in the Ramos V region into unmutated and mutated residues (Fig. S3C). Next, we measured PRO-seq read counts (RPM) in a 7 bp window centered at each C:G pair, that is, C:G ± 3 bp, and selected only non-overlapping windows for analysis (Fig. S3C and Methods). We further divided the mutated group into ten equal sub-groups (deciles) such that sub-group 1 contained the least mutated residues and sub-group 10 contained the most mutated residues. The results, displayed as box plots, showed that there was no discernible increase in PRO-seq density with increasing mutation frequency (Fig. S3D, first row). A Wilcoxon rank sum test following multiple testing correction revealed a lack of statistical significance (defined as *P* < 0.05) between the unmutated and any mutated group, as well as between any pair of mutated groups.

Conversely, we asked whether mutated C:G residues within locales of higher transcription were likely to have higher mutation rates than mutated C:G pairs in locales with lower transcription. To do so, we divided all the 7 bp windows from the mutated group in Fig. S3C into deciles such that sub-group 1 contained the least transcribed windows and sub-group 10 contained the most transcribed windows (Fig. S3E). We then generated boxplots for the mutation frequency of the central C:G residue within the 21 bp windows. The results showed that there was no discernible increase in mutation frequency with increasing PRO-seq density, and no statistical significance (*P* < 0.05 using the Wilcoxon rank sum test) was observed between any pair of sub-groups (Fig. S3F). Collectively, these analyses suggest that mutation frequency is unrelated to local Pol II stalling or local transcription strength.

PRO-seq in AID^−/−^ Eμ^−/−^ cells revealed that nascent sense transcription was uniformly decreased across the V region and intron (Fig. S4A). Of note, PRO-seq showed that the Eμ enhancer harbors the start sites of an antisense transcript, substantially (∼6-7-fold) weaker than the sense transcript (note the Y axis scale in Fig. S4A and quantification in Fig. S4B). This antisense transcript initiates at the 5’ boundary of Eμ and terminates within the V region, resulting in convergent transcription within the V region (Fig. 3 C and D). Importantly, the decrease in antisense transcription in AID^−/−^ Eμ^−/−^ cells (6-10-fold) is considerably stronger than sense transcription (∼2-fold) suggesting that convergent transcription is strongly reduced in the V region and intron (Fig. S4B). Given the weak SHM phenotype in AID^−/−^ Eμ^−/−^ cells (Fig. 2C), this suggests that convergent transcription may not be a major player in V region SHM, in agreement with a recent study wherein SHM on model reporter substrates occurred without substantial antisense transcription ^55^.

We conclude that although some mutations may result from the presence of recurrent stalled Pol II species in their locales, in general, the SHM patterns bear no significant relationship with zones of Pol II stalling, Pol II accumulation or convergent transcription. In addition, SHM is resistant to major decreases in Pol II levels within the V region and intron in both sense and antisense orientations.

### Absence of correlation between Pol II stalling and SHM at two additional human V regions

To determine the transcriptomic profiles of other human V regions, we developed a strategy to replace the endogenous *IGH* V region of Ramos cells with any V region of choice using CRISPR editing (Fig. S5 A and B, and Methods). We chose two V regions corresponding to the unmutated precursors of broadly neutralizing antibodies identified in individuals infected with human immunodeficiency virus 1 (HIV1). One V region consisted of VH4-59*01, DH2-02 and JH6*03 ^92^ and the other of VH3-30*18, DH2-02 and JH6*02 ^93^, which we henceforth refer to as VH4-59 and VH3-30, respectively. In both cases, the sequences upstream and downstream of the new V regions, including the VH4-34 promoter, leader peptide and intron are identical to the endogenous Ramos *IGH* sequence. Sub-regions of both VH4-59 and VH3-30 harbored multimapping reads with the most prominent being the framework region 3 preceding CDR3 in VH4-59 which was entirely covered with multimapping reads (Fig. S5 C and D).

We analyzed the transcriptional and mutational landscape of these two V regions using PRO-seq, PRO-cap and MutPE-seq (Fig. 4A-F). At VH4-59, we observed an increase in PRO-seq enrichment towards the 3’ end of the V region (Fig. 4 A and B), reminiscent of Pol II stalling and akin to that seen at the endogenous Ramos V region (Fig. 3 B and C). Such enrichments were absent within the VH3-30 V region (Fig. 4 D and E). A reasonable explanation is that VH4-59 and VH4-34 belong to the same V gene family (VH4) and hence share higher sequence identity than VH3-30 which is phylogenetically more distant from the VH4 family ^94^. It is also apparent from these analyses that there is no conserved V region transcriptional signature. It is plausible that sequence-intrinsic features may be responsible for the distinct nascent transcription profiles of the different V regions.

**Figure 4:**
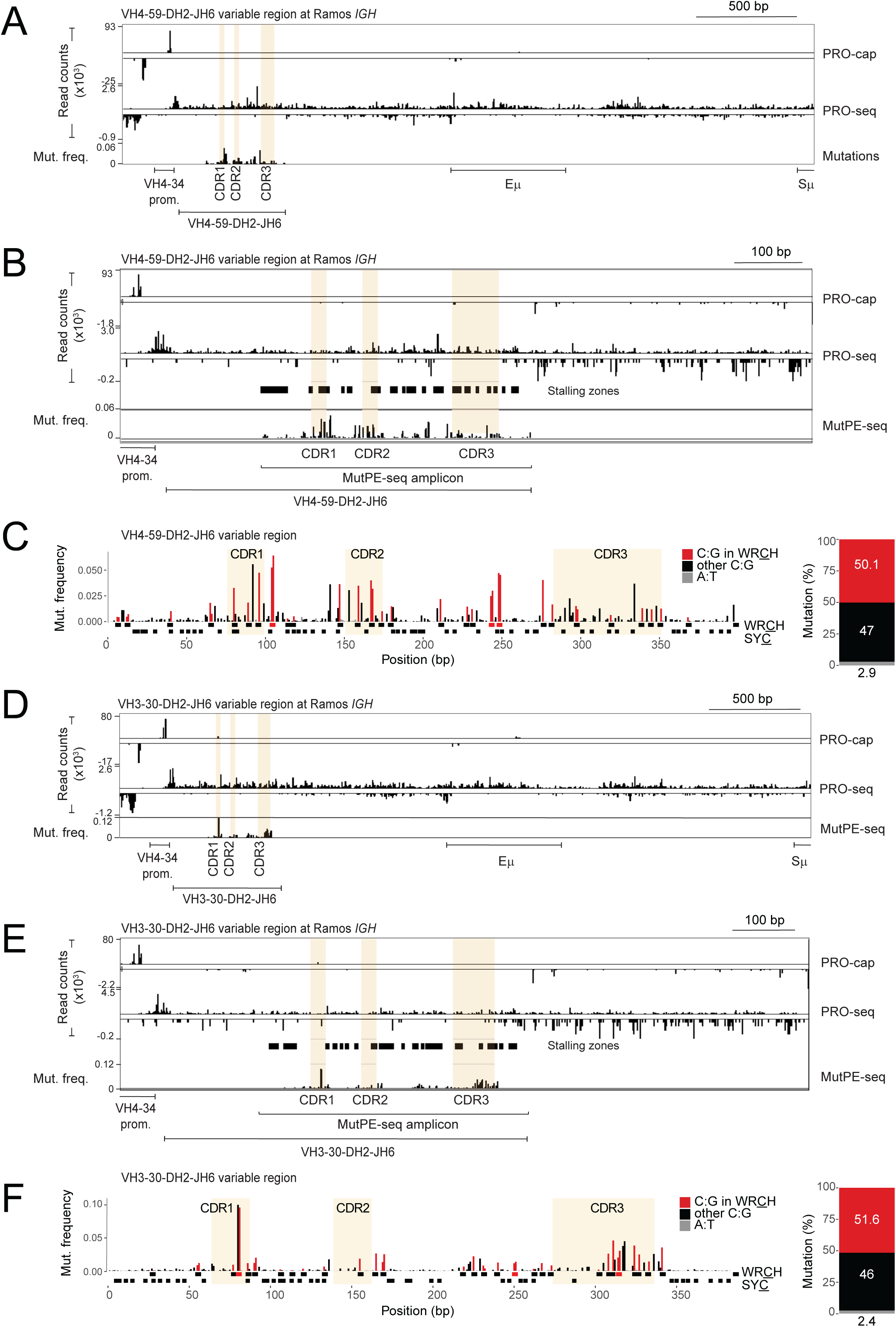
Comparison of transcriptional and mutational landscapes at two different human V regions expressed from the Ramos *IGH* locus. (**A**) IGV browser snapshots of nascent RNA 5’ (PRO-cap) and 3’ (PRO-seq) ends and mutation tracks (MutPE-seq) at the VH4-59-DH2-JH6 V region expressed from the Ramos VH4-34 promoter (CDRs 1-3 highlighted). Mutation analysis was performed following infection of AID^−/−^ cells with AID(JP8Bdel)-expressing lentiviruses for 7 days. **(B)** A magnified view of the VH4-59-DH2-JH6 V region from A above. The stalling zones are shown in the panel between the PRO-seq and MutPE-seq tracks. **(C)** Detailed mutational analysis of the VH4-59-DH2-JH6 V region displayed and color-coded as in Fig. 3E. A bar plot, as in Fig. 3G, with the percentage of mutation frequencies is shown on the right. **(D)** Nascent transcriptional and mutational profiles of the VH3-30-DH2-JH6 V region as in A. **(E)** A magnified view of the VH3-30-DH2-JH6 V region from D above. The stalling zones are shown in the panel between the PRO-seq and MutPE-seq tracks. **(F)** Detailed mutational analysis of the VH3-30-DH2-JH6 V region (see C above for details).

At VH4-59, the location of some highly mutated residues correlated with increased Pol II stalling in their locales, such as in the framework region between CDR2 and CDR3 (Fig. 4B). However, many other similarly mutated residues within and between CDR1 and CDR2 did not show elevated Pol II stalling in their locales (Fig. 4B). Similarly, at VH3-30, the most highly mutated residues located in CDR1 and CDR3 were not associated with Pol II stalling (Fig. 4E). To analyze this systematically, we defined stalling zones in VH4-59 and VH3-30 and compared mutation frequencies between these and non-stalled zones, as described above. For both V regions, there was no significant difference in mutation load between stalled and non-stalled zones (Fig. S3A-B; *P* = 0.6 for VH4-59 and *P* = 0.26 for VH3-30). We also compared mutation frequencies and PRO-seq densities in the neighborhood of C:G pairs at VH4-59 and VH3-30, which revealed that locales with higher rates of mutations did not show a statistically significant correlation (based on the Wilcoxon rank sum test) with transcriptional strength (Fig. S3C-D). Conversely, locales with higher transcription were not more likely to harbor highly mutated C:G residues (Fig. S3E-F).

Of note, when comparing the mutation patterns between all three V regions analyzed here, the strong bias for C:G mutations typical of Ramos cells is seen in all cases (Fig. 3E and Fig. 4 D and F). Moreover, there appears to be some bias of mutation load in the CDRs although several major mutation hotspots are also located within the intervening framework regions (Fig. 3E and Fig. 4 D and F). The coldspot SY<u>C</u> motifs are weakly targeted or untargeted by AID, as expected (Fig. 3 E and Fig. 4, D and F). Finally, the differential mutability of WR<u>C</u>H motifs by AID, ranging from highly mutated to unmutated, is seen in all the analyzed V regions (Fig. 3E and Fig. 4 D and F) and is in line with observations in previous studies ^81,95^.

In sum, the results from VH4-34, VH4-59 and VH3-30 analyses lead us to conclude, firstly, that recurrent Pol II stalling is not present across the entire length of V regions but is restricted to specific sub-regions. Secondly, antisense and convergent transcription is much weaker than sense transcription within V regions. Finally, the data suggest that although Pol II stalling may contribute towards mutagenesis in some locales, in general, the discrete SHM profiles of V regions cannot be explained by recurrent patterns of stalling or Pol II accumulation generally.

### SHM in germinal center B cells at a murine V region shows no direct relationship with nascent transcriptional features

In mice, SHM occurs robustly *in vivo* in germinal center B cells (GCBs) but is inefficient *in vitro* in activated primary B cells ^95^. Thus, we asked if such differences could be related to differences in the underlying transcriptional landscape. To address this, we made use of the B1-8^hi^ *Igh* knock-in mouse where the murine B1-8^hi^ V region is knocked-in at the endogenous *Igh* locus ^96^. In this system, one can compare SHM and transcriptional features of the B1-8^hi^ V region from both germinal center B cells (GCBs) as well as primary *in vitro* activated B cells. To boost SHM in primary cells, we generated homozygous *Igh*^B1-hi/B1–8hi^ *Rosa26*^AIDER/AIDER^ mice where AID fused to the estrogen receptor (AIDER) is expressed constitutively from the *Rosa26* promoter such that upon addition of 4-hydroxytamoxifen (4-HT), AIDER is translocated into the nucleus ^90^.

PRO-seq analysis from homozygous B1-8^hi^ mice showed that the VH1-72 gene used in the B1-8^hi^ recombined V region shares strong homology with other V genes in the murine *Igh* locus resulting in a very high degree of multimapping (Fig. S5E). Importantly, the CDR3 region was covered with uniquely mapping reads since this region is formed via the junction of the V, D and J segments during VDJ recombination resulting in a unique sequence in the genome (Fig. S5E). PRO-seq and PRO-cap showed that there were no striking differences at the B1-8^hi^ V region between GCBs and primary B cells activated with LPS, IL-4 and RP105 (Fig. 5A). Moreover, we observed local zones of Pol II accumulation, internal initiation events and antisense/convergent transcription, but these did not necessarily correlate with zones of mutation (Fig. 5A).

**Figure 5:**
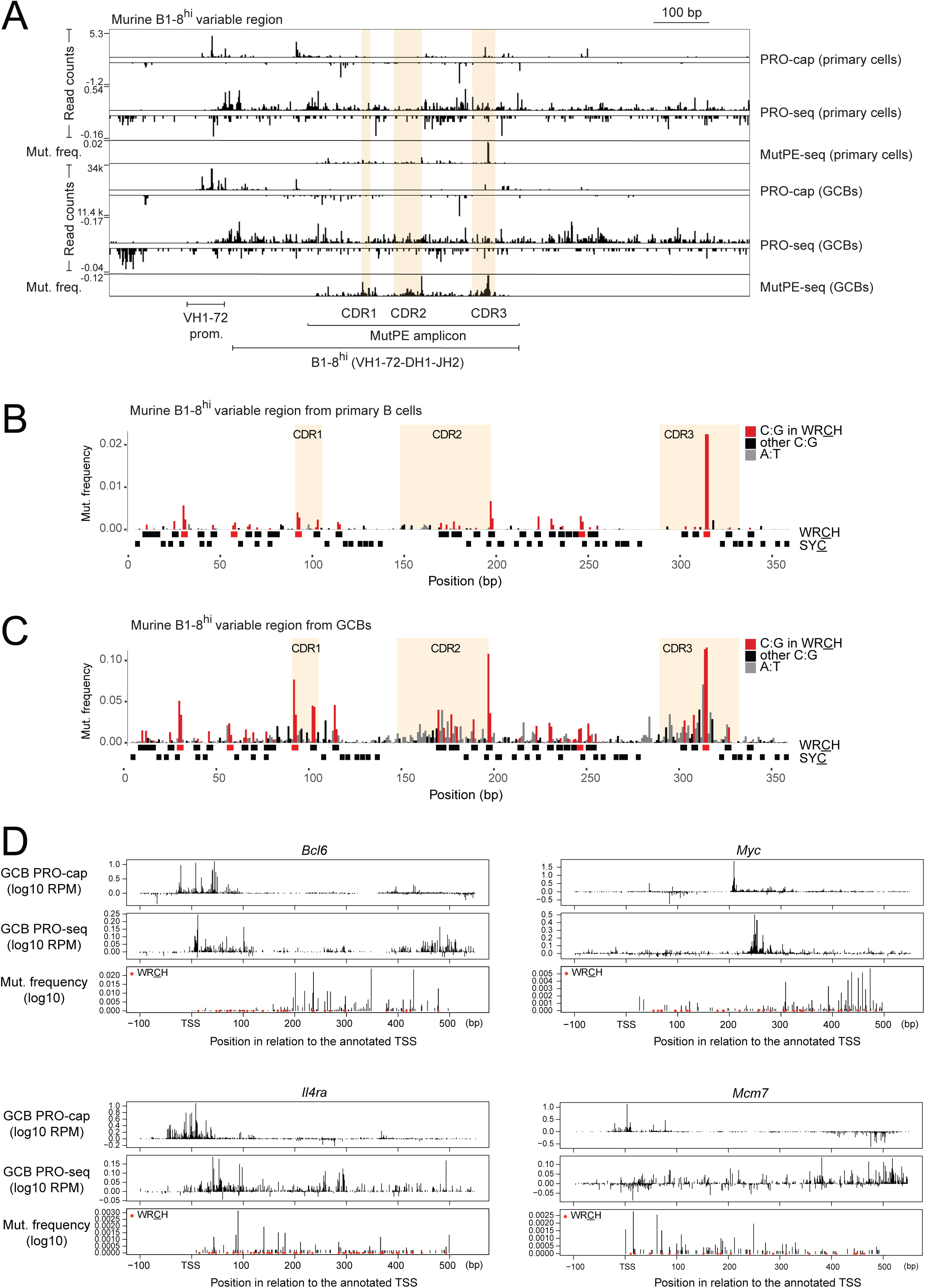
Transcriptional and mutational landscapes of the murine B1-8^hi^ V region and non-*Ig* AID target genes in mice. **(A)** Nascent transcriptional analysis at the *Igh* locus in murine B1-8^hi^ primary, splenic B cells stimulated for four days with LPS, IL4 and RP105. IGV browser snapshots show the 5’ (PRO-cap) and 3’ (PRO-seq) ends of the aligned reads. For MutPE-seq, *Igh*^B1–8hi/ B1–8hi^ *Rosa26*^AIDER/AIDER^ primary B cells were activated with LPS, IL4 and RP105 for 4 days with 4-HT. PRO-seq, PRO-cap and MutPE-seq were also performed from sorted, splenic germinal center B cells (GCBs) following immunization with sheep red blood cells for 10 days. **(B)** Analysis of SHM at the B1-8^hi^ V region from B1-8^hi^ *Rosa26*^AIDER^ primary, activated murine splenocytes. The bar graph on the right shows the percentage of each indicated mutation category (see Fig. 3E and 3G for details). **(C)** Analysis of SHM at the B1-8^hi^ V region from splenic GCBs following immunization with sheep red blood cells for 10 days. **(D)** 5’ and 3’ ends of nascent RNA (PRO-cap, PRO-seq) and mutation profiles at four selected AID target genes. Mutational data are from Alvarez-Prado et al. (2018) wherein the first 500 bp of the genes were sequenced. The region displayed extends from –100 bp from the annotated (RefSeq) TSS up to 50 bp downstream of the sequenced amplicon. The WR<u>C</u>H motifs are indicated as red dots.

We note that the PRO-seq read densities at the B1-8^hi^ V region from GCBs was substantially lower than those seen at the V regions analyzed in Ramos cells. We attribute this to the fact that GCBs are more fragile than primary cells or cell lines resulting in considerable loss of viability and quality during cell sorting. Consequently, the efficiency of the run-on reaction from isolated GCB nuclei, the first step of PRO-seq, is much less efficient resulting in lower numbers of aligned reads in the final libraries. These weak signals precluded a robust identification of stalling sites and hence we were unable to perform the comparative analysis of mutation frequency and stalling zones as we did for the other V regions described earlier.

As noted previously ^95^, SHM profiles of B1-8^hi^ GCBs are not identical to primary B cells because GCBs exhibit 5-10 fold higher frequencies of SHM and substantial mutagenesis of A and T residues arising from error-prone DNA repair of AID-induced mismatches at neighboring C:G residues (Fig. 5 B and C). The palindromic AGCT hotspot in CDR3 is selectively mutated with high frequency in primary cells, in line with a former report showing that this hotspot is the earliest targeted motif both *in vitro* and *in vivo* ^95^. Importantly, however, closer inspection showed that even in primary B cells, the same WR<u>C</u>H motif appears to be mutated as in GCBs suggesting that AID targeting specificity is not substantially different between primary B cells and GCBs (Fig. 5 B and C), in line with their comparable transcriptional landscapes (Fig. 5A). AID activity is linked to cell division ^97,98^ and in our experiments, GCBs were harvested after 10 days of immunization whereas primary cells were harvested after 4 days of 4-HT treatment. Hence, it is plausible that GCBs may have acquired higher rates of mutation, in part, due to more rounds of cell division within germinal centers compared to primary cultures.

We conclude that the differences in mutation frequency between primary B cells and GCBs cannot be explained by underlying nascent transcriptional features. Consequently, as in the case of human V regions described above, the nascent transcriptional landscape of the B1-8^hi^ V region is not predictive of the patterns or frequencies of SHM.

### Analysis of non-IG AID target loci reveals no correlation between SHM and transcription strength

To extend our analysis to non-IG AID targets, we made use of a previously available SHM dataset from murine GCBs where mutation frequencies at 275 AID target genes were identified by deep sequencing the first 500 bp from the annotated transcription start site (TSS) ^17^. In this study, mutation analysis was performed in GCBs from mice deficient in base excision and mismatch repair pathways (*Ung*^−/−^*Msh2*^−/−^). In these mice, the processing of AID-induced U:G mismatches is abolished and, consequently, DNA replication over these lesions leads to C to T and G to A transition mutations that represent the direct targets of AID ^99^. By comparing these data with PRO-seq and PRO-cap obtained from GCBs, we could compare mutation profiles with those of nascent transcription. Importantly, at many genes, initiation sites or zones defined by PRO-cap are often located at considerable distances from the annotated TSS (defined by the RefSeq database and labeled as TSS in Fig. 5D, Fig. S6 and Fig. S7). Consequently, the 500 bp segment sequenced for mutational analysis ^17^ often begins further upstream or downstream of the initiation sites newly defined by PRO-cap.

The large number of genes in this analysis led to new and unexpected observations regarding the nature of AID targeting. At virtually all genes, the differential mutation of WR<u>C</u>H motifs was evident in that many such motifs were mutated to varying degrees with many others being unmutated. Moreover, several non-WR<u>C</u>H cytidines were as efficiently mutated as WR<u>C</u>H cytidines (Fig. 5D and several additional examples in Fig. S6 and Fig. S7). In V and switch regions, mutations initiate ∼100-150 bp after the promoter suggesting that AID may be excluded from initiating or early elongating Pol II ^2,3,62,100,101^. However, we identified a set of genes where the highest mutation frequencies were observed within 100-150 bp of the initiation site defined by PRO-cap (*Il4ra* in Fig. 5D and additional examples in Fig. S6A). Indeed, in some of these genes, the most mutated residues were in very close proximity to the initiation site, such as *Mcm7* (Fig. 5G). In some other genes, strongly mutated residues coincided with the strongest peak of initiation (Fig. S6B). These observations indicate that AID can act at the early stages of the transcription cycle at least at some genes, suggesting that AID is not *per se* excluded from initiating or early elongating Pol II.

As we observed in V regions, mutation frequencies at individual nucleotides and mutation zones did not necessarily correlate with enrichments of PRO-seq or PRO-cap (Fig. 5D, Fig. S6 and Fig. S7). For example, at *Bcl6*, the major zone of mutation lies in a region of low PRO-seq signal where many mutations are not in WR<u>C</u>H motifs (Fig. 5D). At *Myc*, mutations are observed upstream of the initiation site defined by PRO-cap indicating that mutagenesis can also occur in the antisense orientation (Fig. 5E). Thus, as at V genes, SHM does not correlate with local nascent transcriptional features.

To systematically determine if mutation frequency corelated with local transcriptional strength, we divided all C:G residues from Alvarez-Prado et al.^17^ into three groups. Group 1 contained C:G residues in unmutated genes, group 2 contained unmutated C:G within the mutated genes and group 3 contained the mutated C:G residues (Fig. S8A). The C:G residues in group 3 were ranked from lowest to highest mutation frequency and further divided into ten equal groups (deciles) where residues in sub-group 1 were the least mutated and those in sub-group 10 were the most mutated. Next, we counted PRO-seq reads in a 7 bp window centered at each C:G residue (C:G ± 3 bp) (Fig. S8A) and plotted the results as box plots (Fig. S8B).

There was no visible trend of higher PRO-seq density in groups with higher mutation frequency although statistical analyses showed that the PRO-seq densities in group 1 were significantly lower than most, but not all, other groups, in line with the fact that mutation rates are higher in more transcribed genes ^17,24^ (Fig. S8B). However, the difference in mean PRO-seq values between group 1 and other groups ranged between 1.2-1.9-fold. Based on our results from Eμ^−/−^ cells where >2-fold differences did not have a major impact on SHM (Fig. 2), we infer that the reason for the absence of mutation in group 1 is not solely due to lower transcription.

Group 2 showed a significant difference in PRO-seq density only with sub-group 8 within group 3. Moreover, the PRO-seq densities in group 3, sub-group 10, the most mutated group, were actually lower than in many other groups and even showed statistically significant differences in some cases (Fig. S8B). This suggests that, within AID target genes, highly mutated C:G residues are not generally located in genomic neighborhoods harboring higher levels of transcription and Pol II occupancy.

Conversely, we asked whether C:G residues within locales of higher transcription were likely to have higher mutation rates. As was done previously for V regions (Fig. S3E-F), we divided all the 7 bp windows from group 3 above into deciles such that sub-group 1 contained the least transcribed and sub-group 10 contained the most transcribed windows, respectively (Fig. S8C). We then generated boxplots for the mutation frequency of the central C:G residue within the 7 bp windows (Fig. S8 C and D). The results showed no trend of higher mutation frequency in groups with higher PRO-seq values with only one pair (group 3 versus group 10) showing a statistically significant difference (Fig. S8D). We conclude that the level of local transcription within mutated genes is not predictive of C:G mutability.

### SHM targeting of the murine B1-8^hi^ V region is retained in the context of the human *IGH* locus

To determine whether the gene regulatory context affects the patterns of SHM and transcription, we replaced the endogenous V region in Ramos cells with the entire murine B1-8^hi^ V region (Ramos^B1–8hi^) using the strategy described earlier (Fig. 6A and Fig. S4, A and B). In this scenario, transcription and SHM of the murine B1-8^hi^ V region would be under the control of the Ramos VH4-34 promoter and the human Eμ and 3’ *IGH* super-enhancers. This allowed us to ask whether the targeting of SHM to the B1-8^hi^ V region was influenced by differences in the human versus the mouse enhancers and promoters. Importantly, when placed in the context of the human *IGH* locus, the B1-8^hi^ sequence was covered almost exclusively by uniquely mapping reads (Fig. 6B), which is in sharp contrast to the extensive multimapping observed within B1-8^hi^ at the murine *Igh* locus (Fig. S4E). This is because the VH1-72 gene used in the B1-8^hi^ sequence is much less homologous with the human V genes than the mouse V genes.

**Figure 6:**
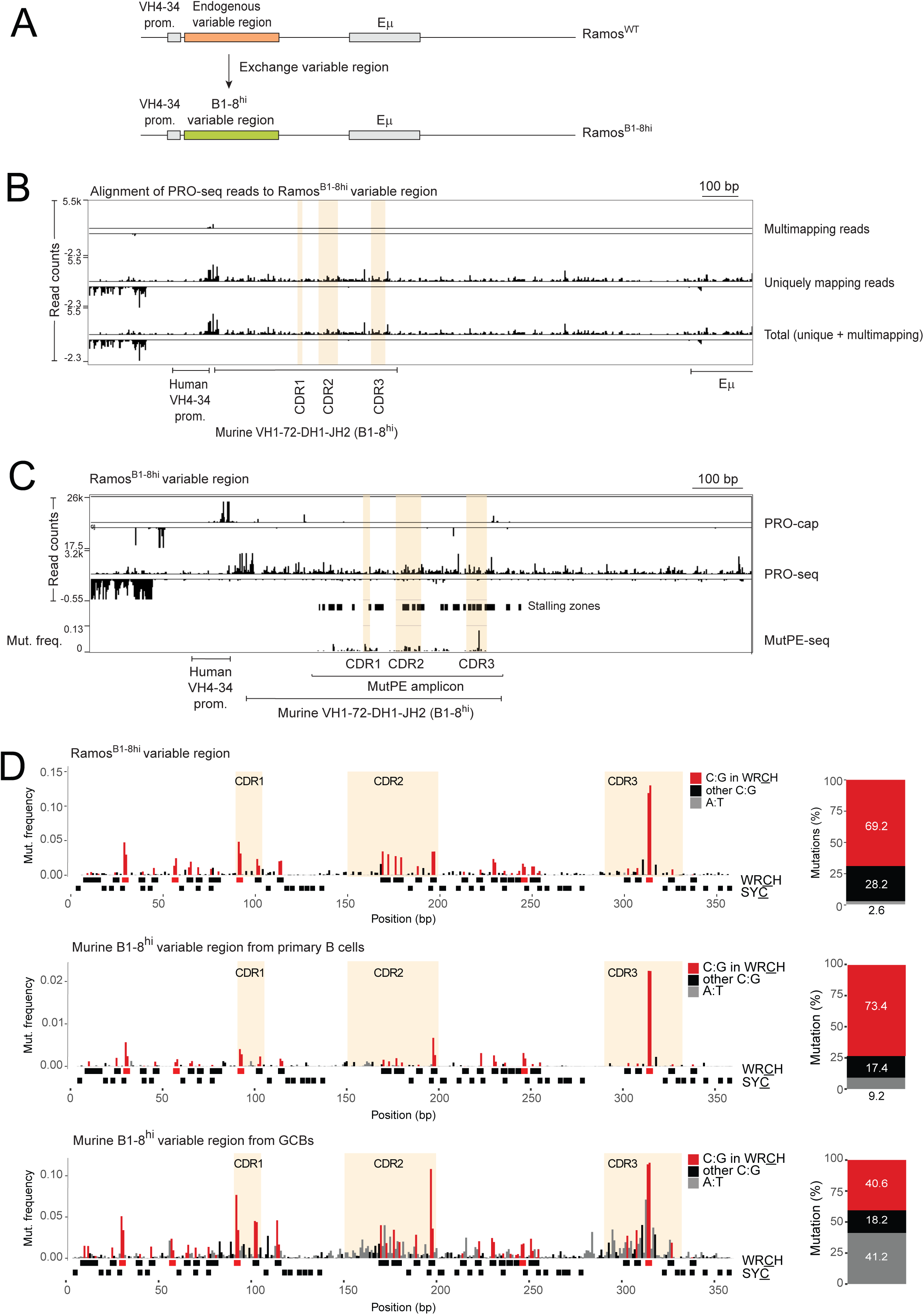
Analysis of gene regulatory context on the nascent transcriptional landscape and SHM of the murine B1-8^hi^ V region. **(A)** Scheme showing the exchange of the endogenous Ramos V region for the murine B1-8^hi^ V region to generate the Ramos^B1–8hi^ human *IGH* locus following the approach described in Fig. S2A. **(B)** 3’ ends of nascent RNAs by PRO-seq at the Ramos^B1–8hi^ *IGH* locus showing the distribution of multimapping, unique and total signal. **(C)** Nascent RNA 5’ and 3’ ends of PRO-cap and PRO-seq respectively and MutPE-seq analysis at the Ramos^B1–8hi^ *IGH* locus. MutPE-seq was performed following infection with AID(JP8Bdel)-expressing lentiviruses for 7 days. The stalling zones are shown in the panel between the PRO-seq and MutPE-seq tracks. **(D)** Details of the mutation patterns of the B1-8^hi^ V region expressed from the Ramos *IGH* locus obtained by MutPE-seq. The bar graphs on the right show the percentage of each indicated mutation category (see Fig. 3E and 3G for details). For comparison, the murine B1-8^hi^ mutation profile from primary murine B cells (middle panel) and murine GCBs (bottom panel), exactly as in Fig. 5B-C, are included.

PRO-seq and PRO-cap revealed that the nascent transcriptional profile of the B1-8^hi^ sequence in Ramos^B1–8hi^ cells (Fig. 6C) shared similarities with the B1-8^hi^ sequence in the murine *Igh* context (Fig. 5A). For example, similarities were noted in CDR3 and flanking sequences, for instance, the presence of the antisense initiation site (PRO-cap track) and shared spikes of nascent transcriptional activity (PRO-seq) (Fig. 6C). More importantly, MutPE-seq following AID (JP8Bdel) infection revealed a very similar SHM pattern in Ramos^B1–8hi^ cells compared to the murine B1-8^hi^ context, especially with primary murine B cells (Fig. 6D). Of note, the dominant, palindromic AGCT hotspot in CDR3 was the most mutated residue exactly as in B1-8^hi^ murine primary cells and GCBs (Fig. 6D). The other mutated C:G residues in the Ramos^B1–8hi^ sequence were largely the same as those mutated in the murine context albeit the relative mutation rates within each V region varied (Fig. 6D, compare with Fig. 5 B and C). Stalled Pol II zones within the Ramos^B1–8hi^ V region bore no significant relationship with mutation frequency compared to non-stalled zones (Fig. S3A-B; *P* = 0.83). Analysis of PRO-seq density in the neighborhood of mutated and unmutated C:G pairs revealed a lack of statistical significance between mutation frequency and local transcriptional strength (Fig. S3C-F). Thus, as in the case of the other V regions analyzed here, there is no significant relationship between Pol II stalling, Pol II occupancy and local SHM at the Ramos^B1–8hi^ V region.

We note that mutation frequencies in murine GCBs and AID (JP8Bdel)-expressing Ramos cells were comparable (note the Y axes scales in Fig. 6D) with the major difference being the weak A:T mutagenesis typical of Ramos cells. Of note, the sense PRO-cap peak in CDR3 in the murine B1-8^hi^ (Fig. 5A), located just 3 bp upstream of the strong AGCT hotspot, is absent in Ramos^B1–8hi^ (Fig. 6C) implying that this initiation site is not essential for the high mutation frequency of this hotspot motif. In sum, the nascent transcriptional and SHM patterns of the B1-8^hi^ V region in mouse B cells are largely retained in Ramos^B1–8hi^ cells.

Collectively, our results lead us to conclude that, following AID recruitment via IG enhancers and its association with Pol II-associated factors, the discrete and reproducible mutation patterns in V regions and non-Ig SHM target genes are apparently “hard-wired” into the DNA sequence rather than the result of the local Pol II stalling, convergent transcription or activating chromatin marks.

## Discussion

In this study, we explored the relationship between transcription and V region SHM by performing a detailed comparative analysis of SHM with nascent transcription and activating chromatin marks.

Our analysis of the epigenetic landscape showed that major decreases in activating chromatin marks and nascent transcription, via deletion of the Eμ enhancer, do not have a significant impact on SHM. Although this does not rule out a role for these marks in creating an optimal chromatin environment for SHM, it does suggest that the absolute levels of these marks do not correlate with SHM. Intriguingly, compared to other highly active genes, including non-Ig AID targets ^24,102^, the V region chromatin locale is unusual for two reasons. First, it does not harbor the typical bimodal distribution of H3K27ac and H3K4me3 at the TSS despite robust bidirectional transcription and normal nucleosome occupancy as measured by the levels of histone H3. Second, the V gene body is marked by relatively low levels of these marks. However, it remains unclear whether this locale contributes to making the V genes among the most favored targets of AID in B cells.

The key conclusion of our study is that local patterns and frequencies of SHM, either in V regions or non-Ig AID targets, cannot be predicted from, nor do they appear to be derived from, sites of Pol II stalling, internal initiation or convergent transcription. Our approach clearly detected all these features within all four V regions analyzed here, in line with the global gene-level correlations previously noted ^17,27,51^. However, by accurately mapping the precise locations of these features within V regions and non-IG SHM target loci, our study revealed that at the local sequence level, the presence of these features did not generally correlate with increased local SHM. Indeed, several zones of elevated SHM in different V regions and non-Ig AID target loci appeared to be entirely devoid of any discernible transcriptional features. This implies that these features may not be major players in SHM and, therefore, that the ssDNA associated with them may not be the major substrate for AID. Moreover, the nascent transcription landscapes of the four V genes that we report in this study are quite dissimilar, which argues against any conserved V region nascent transcriptional signature. Finally, the murine B1-8^hi^ V region retains its characteristic SHM profile *in vitro* and *in vivo*, and in both murine and human *cis*-regulatory contexts. Altogether, these results suggest that the discrete SHM patterns of individual V regions may be influenced by local sequence context, rather than specific local transcriptional features.

### What is the role of transcription in SHM?

The consistent lack of correlation between transcriptional features and SHM reported here begs the question of what role transcription plays in SHM. SHM depends on the availability of ssDNA, but the source(s) of accessible ssDNA for AID remains a major unresolved question. It has been proposed that ssDNA within transcribing Pol II complexes could be a substrate for AID, especially when Pol II is paused or stalled, an event that would lead to increased ssDNA exposure ^17,27,52–56^. However, we found no significant correlation between SHM frequency and Pol II stalling zones or Pol II accumulation, which prompted us to take a closer look at the structures of paused and elongating Pol II complexes. This analysis showed that of the ∼11 nucleotide non-template ssDNA in the structure, about half lies buried within Pol II ^88,89^ (Fig. 7A). Importantly, the exposed portion (∼5 nt) is entirely covered by SPT5 ^89^ (Fig. 7A). The PAF complex and SPT6 are located further away from this site ^88^ (Fig. 7A). Under such structural constraints, it is difficult to conceive how AID could access the ∼5 nt exposed ssDNA and catalyze deamination even if Pol II were paused or stalled. Thus, it appears unlikely that the transcription bubble within an actively elongating Pol II holoenzyme, including a stalled complex, serves as an optimal substrate for AID.

**Figure 7:**
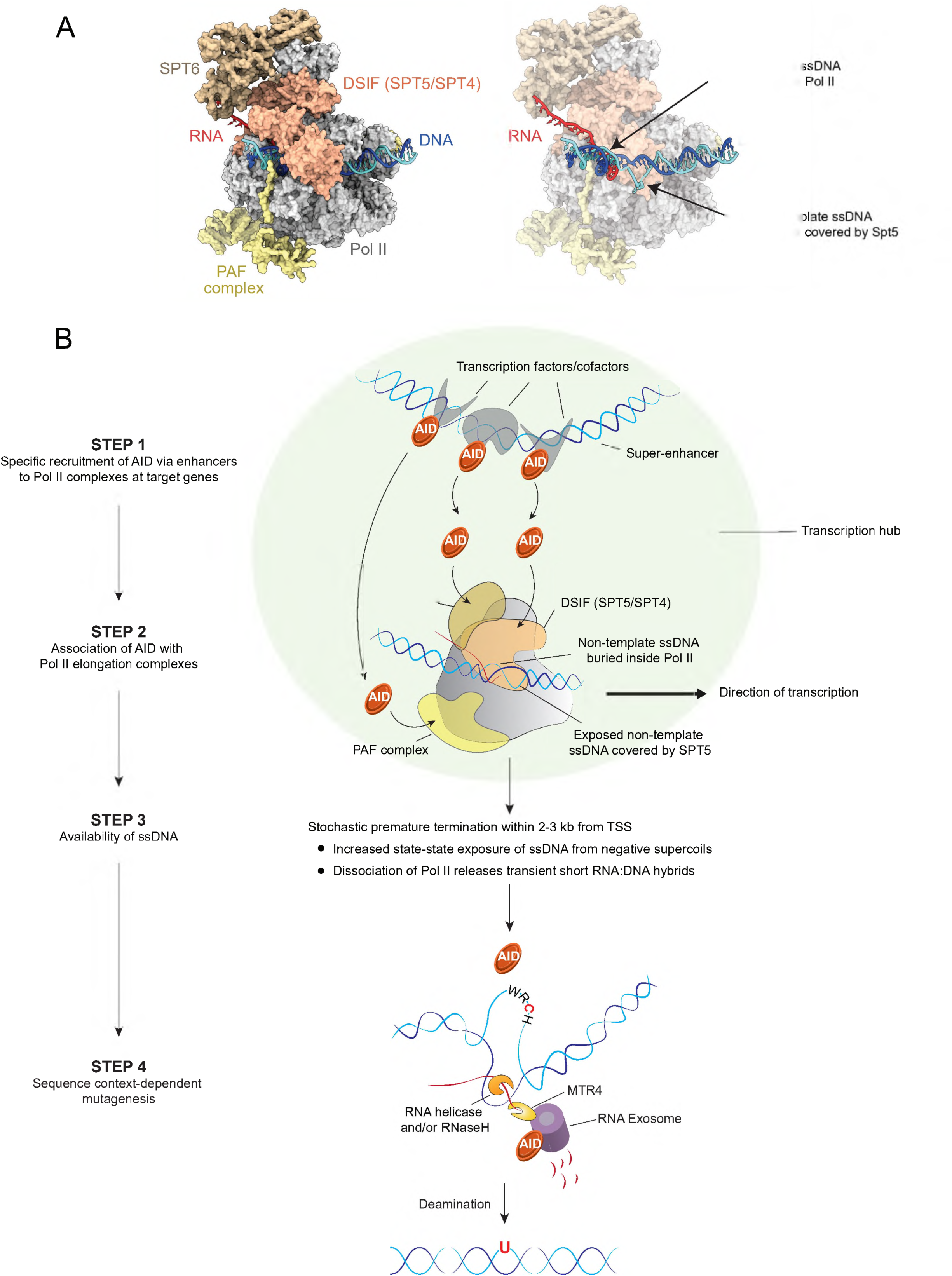
Integrative model for co-transcriptional AID targeting and differential motif mutability. (**A**) 3D structure of the human RNA Pol II elongation complex visualized with ChimeraX. We superimposed, via their RPB1 subunits, the elongation complex structure (PDB ID 6GMH) onto the transcribing Pol II-DSIF (a complex of SPT5 and SPT4) structure (PDB ID 5OIK). Proteins are shown as surfaces (Pol II, grey; DSIF, salmon; SPT6, brown; PAF complex, yellow) and nucleic acids as cartoons (DNA template strand, blue; DNA non-template strand, cyan; RNA, red). The right panel highlights the trajectory of DNA and RNA buried within Pol II. The exposed non-template strand of the transcription bubble is occluded from interactions with AID due to it being completely covered by SPT5. **(B)** Proposed model for SHM that incorporates various known aspects of SHM biology. The cartoon diagram of the Pol II elongation complex reflects the actual structure described in (A) above and in the main text. AID is recruited to the Pol II complex via super-enhancers in the context of a transcriptional hub (step 1) and is then retained via interactions with elongation factors, SPT5, PAF and SPT6 (step 2). The next step is the availability of ssDNA (step 3). We support the idea that premature transcription termination may be an important source of ssDNA ^48^. This event is not associated with stalling but occurs randomly within the first 2-3 kb from the TSSs ^106^. A slower-moving terminating Pol II may provide a more stable source of upstream, negative supercoiled DNA favoring higher AID activity. Upon dissociation of Pol II, the non-template strand is likely available as ssDNA for some time since the DNA:RNA hybrid would prevent immediate reannealing. The RNA is removed by RNA helicases and RNaseH followed by degradation of the RNA by the RNA exosome, which is known to associate with AID and provide access to the template strand. In this manner, AID can access ssDNA through various means all in the context of termination.

Nevertheless, various studies have provided important insights into potential alternative sources of ssDNA. For instance, supercoiled DNA, but not relaxed DNA, can serve as efficient substrates for AID *in vitro* in biochemical assays and *in vivo* in bacterial SHM assays in the complete absence of transcription ^44,45,57^. In fact, untranscribed supercoiled plasmids can be mutated *in vitro* at rates comparable to those seen in V regions *in vivo* ^45^. The proposition is that negatively supercoiled DNA, which occurs in the wake of transcribing Pol II and is inherently prone to transient melting, can generate ssDNA patches on either strand as substrates for AID ^44,45,57^. Indeed, transcription-dependent ssDNA patches have been detected in SHM model substrates, V regions and other AID target loci ^54,55,57,58^. Importantly, however, these studies revealed that the location and frequency of ssDNA patches did not correlate with the patterns and spectra of SHM, implying that the incidence of ssDNA as detected by these assays is not a predictor of SHM patterns ^55,57,59^.

Another key insight is the fact that significant mutational asymmetry is observed upon depletion of RNA Exosome subunits or the DNA:RNA helicases, Mtr4, Senataxin and others ^103–105^, which implies that one of the major species targeted by AID is an RNA:DNA hybrid. Indeed, the V region sense transcript is a substrate of the RNA exosome and the latter is required for optimal mutation of the template strand in the B1-8^hi^ V region, findings that directly implicate the processing of DNA:RNA hybrids, or short transient R loops, in directing the patterns of SHM ^103^.

Nevertheless, because all the above-mentioned features are continuously associated with transcription, they cannot, by themselves, explain the distinct and highly reproducible SHM profiles of V regions, nor can they explain why SHM spectra are restricted to a ∼2 kb window starting from the V gene promoter ^3,62,63^. To reconcile these observations, we support a model centered on premature transcription termination, which, as proposed below, can define the boundaries of SHM and also provide a source of ssDNA, the latter having been suggested previously in the context of RNA exosome-mediated AID targeting ^49,50^ as well as SHM on plasmid substrates and in the Ramos V region ^47,48^ (Fig. 7B).

It is now appreciated that pervasive termination occurs near the 5’ ends of all active genes and enhancers resulting from the activity of the Integrator endonuclease complex ^106–108^. Specifically, the depletion of Integrator subunits manifests as an increase in Pol II and nascent transcript levels within the first 3-4 kb from TSSs ^106–108^, which, intriguingly, encompasses the known spectrum of SHM at V regions, that is, ∼2 kb from the V gene promoters ^3,62,63^. A similar process would be expected to occur at the 5’ ends of non-IG AID target genes or at divergently transcribed enhancers that are targets of AID ^16,55,109,110^. Importantly, we note that accumulation of Pol II in V regions that we (this study) or others ^48,54^ have detected cannot be inferred as termination zones because the ChIP and nascent transcript profiling assays used therein do not provide information on the fate of the detected Pol II species. Studies to date reveal no specific locations for premature termination, rather suggesting that such events are stochastic and an inherent feature of early elongation ^106–108^.

Another attractive aspect of premature termination in the context of SHM is that it can potentially provide multiple avenues for AID to access ssDNA ^47–50^. Firstly, the speed of a terminating Pol II is expected to be lower than a rapidly elongating one, which would increase the steady-state availability of upstream, negatively supercoiled DNA for AID to access. Secondly, although the structural details of how Pol II terminates and dissociates off DNA have not been resolved, it is conceptually plausible that the collapse of the elongation complex could release ssDNA on the non-template strand while the nascent RNA is still hybridized to the template strand, which provides another source of ssDNA (Fig. 7B, step 4). Finally, the RNA would have to be unwound by DNA:RNA helicases ^105^ and/or digested with RNaseH followed by degradation from the 5’ end by Xrn2 and from the 3’ end by the RNA exosome, thereby providing AID access to the template strand ^31,103–105,111^ (Fig. 7B, step 4). Given that IG loci are amongst the most highly expressed in activated B cells, the combined effect of having multiple rounds of transcription per cell cycle coupled with stochastic termination within the first 2-3 kb from V gene promoters can plausibly explain the empirically observed boundary of SHM, that is, ∼2 kb from V gene promoters ^3,62,63^. The model can also explain why, at some non-IG SHM target loci, we observed mutations overlapping with, or in close proximity to, the transcription initiation zone. Most importantly, such stochastic events will not be detected by standard population-based approaches, which can explain the lack of correlation between local SHM frequency and any transcriptional feature, be it Pol II stalling, internal initiation, convergent transcription or ssDNA patches.

Although SHM-specific enhancers could explain why AID does not target all transcribed genes ^25,55^ and the termination model could explain the spectrum of SHM, they do not explain why WR<u>C</u>H motifs are differentially mutated. Indeed, the same motif can show high variation in mutation frequency depending on its location within a given V region ^81,95,112,113^, a feature that is also evident in the V regions and non-Ig genes analyzed in this study. This implies that, beyond AID recruitment and ssDNA availability, the DNA sequence context may determine whether ssDNA containing a WR<u>C</u>H motif is mutated and to what extent. This notion is supported by a biochemical study of SHM on reporter cassettes^114^, *in vivo* analyses of SHM in V regions from mice^81,95^, computational modeling studies of V regions ^112,113,115^ and by altering the nucleotide context of AGCT hotpot motifs *in vivo* and *in vitro* which suggested that a sequence locale enriched in pyrimidine dimers can promote AGCT mutagenesis^116^. In this regard, our study is important in showing that the molecular basis of this sequence context dependency of SHM does not lie at the level of Pol II stalling or any other nascent transcriptional feature, which was one of the initial hypotheses that motivated the present study. In sum, further work is necessary to determine whether sequence context regulates SHM patterns *in vivo* and what the underlying mechanism(s) may be.

## Methods

### Cell culturing

Ramos were cultured in complete RPMI medium (in-house) supplemented with 10% fetal bovine serum (FBS; Invitrogen), glutamine (Invitrogen), sodium pyruvate (Invitrogen), HEPES (made in-house) and antibiotic/antimycotic (Invitrogen). LentiX packaging cells were cultured in complete DMEM medium (in-house) containing 10% fetal bovine serum (FBS; Invitrogen), glutamine (Invitrogen), sodium pyruvate (Invitrogen), HEPES (made in-house) and antibiotic/antimycotic (Invitrogen).

### Mice

The mice were maintained in a C57BL/6 background and housed in the IMBA-IMP animal facility in standard IVC cages with HEPA filtering. B1-8^hi^ Rosa26 ^AIDER^ mice were generated by crossing B1-8^hi^ mice ^80^ with *Rosa26^AIDER^* mice ^90^ and maintained as a homozygous line for both alleles. All animal experiments were carried out with valid breeding and experimental licenses (GZ: MA58-320337-2019-9, GZ: 925665/2013/20 and GZ: 618046/2018/14) obtained from the Austrian Veterinary Authorities and in compliance with IMP-IMBA animal house regulations.

### Generation of AID^−/−^ Ramos cells

We generated Ramos cells expressing EcoR, the receptor for the ecotropic envelope protein, Eco-Env, so as to allow efficient infection of Ramos cells with ecotropic lentiviruses. Ramos cells were infected with a pRRL lentiviral vector expressing EcoR and maintained under Puromycin selection. These cells were transfected with a small guide RNA targeting *AICDA*, the gene expressing AID, and recombinant Cas9 protein via electroporation using the Neon transfection system (Thermo Fisher Scientific). The following day, single cells were sorted into 96-well plates using BD Aria III FACS sorter and allowed to expand. Four weeks later, clones harboring frameshifting indels at the Cas9 target site were identified by genotyping and loss of AID in these clones was confirmed by Western blot analysis. One clone (D3) was used for all subsequent analyses and for generation of new lines.

### Generation of Eμ^−/−^ Ramos cells

To delete the 583 bp segment containing Eμ, we generated homology repair plasmids having Ef1a promoter-driven floxed GFP and mCherry expression cassettes flanked by homology arms. These plasmids were transfected into AID^−/−^ Ramos cells (clone D3) along with three in vitro synthesized guide RNAs (6 μg each, designed using CRISPOR) and 7.5 μg recombinant Cas9 protein (Vienna Biocenter Core Facilities) using the Neon Transfection System (Thermo Fisher Scientific). A week later, GFP/mCherry double-positive single cells were isolated using a BD FACS Aria III sorter (BD Biosciences). Successful knock-in clones were identified by genotyping with PCR and Sanger sequencing of PCR products. Next, 200 μg recombinant Cre recombinase (Molecular Biology Service, IMP) was added to the culture medium to excise the floxed mCherry/GFP cassettes. GFP/mCherry double-negative clones were isolated one week after electroporation using a BD FACS Aria III sorter and genotyped via PCR and Sanger sequencing of PCR products.

### Generation of Ramos cells expressing new, exogenous V regions

AID^−/−^ Ramos cells (clone D3) were electroporated with Cas9-sgRNA ribonucleoprotein complexes (prepared in-house by the Vienna Biocenter Core Facilities – ProTech) and homology repair templates containing a pair of unique sgRNA-target sites to excise the entire V region including the promoter (Fig. S2A). Single, IgM-negative clones were isolated via flow cytometry using a BD Aria III sorter, expanded and genotyped. One validated IgM-negative clone was selected as the parental clone. This clone was then electroporated with Cas9-sgRNA ribonucleoprotein complexes targeting the unique sgRNA sites and homology repair templates containing new V regions under the control of the endogenous Ramos VH4-34 promoter. Single, IgM-positive cells were isolated, expanded and genotyped.

### Lentiviral infections

Lentiviral pRRL vectors expressing AID(JP8Bdel) or AID (m7.3) coupled with mCherry were transfected along with ecotropic envelope (Eco-env)-expressing helper plasmid into LentiX cells via the standard calcium phosphate methodology. Lentiviral supernatants were used to infect Ramos cells via spinfection (2350 rpm for 90 min) in the presence of 8 μg/ml Polybrene (Sigma). Flow cytometry (BD Aria III sorter) was used to sort mCherry-positive cells for mutation analysis.

### Isolation and activation of murine primary, splenic B cells

Mature, naïve B cells were isolated from spleens of 2-4-month-old B1-8^hi^ Rosa26 ^AIDER^ mice, as per established protocols ^27^. B cells were cultured in complete RPMI medium supplemented with 10% fetal bovine serum (FBS) and antibiotics, Interleukin 4 (IL4; made in-house by the Molecular Biology Service, IMP), 25 μg/ml Lipopolysaccharide (Sigma) and RP105 (made in-house by the Molecular Biology Service, IMP) and harvested after 3 or 4 days. For SHM assays from activated B1-8^hi^ *Rosa26* ^AIDER^ primary B cells, 2μM 4-hydroxy tamoxifen (4-HT) (Sigma) was added at the time of activation with IL-4, LPS and RP105.

### Isolation of germinal center B cells (GCBs) from immunized B1-8^hi^ mice

Sheep red blood cells (SRBCs) were washed thrice with PBS of which 0.2×10^9^ SRBCs in 100µl PBS (per mouse) were injected into B1-8^hi^ mice followed by another injection of 1×10^9^ SRBCs in 100µl PBS five days later. Mice were harvested 10 days after the first immunization. Spleens were harvested and single-cell suspensions were stained with B220 conjugated to fluorescein isothiocyanate (B220-FITC, BD Biosciences, 1:500 dilution), Fas conjugated to PE-cyanine 7 (Fas-PE-Cy7, BD Biosciences, 1:1000 dilution) and CD38 conjugated to Allophycocyanin (CD38-APC, ThermoFisher, 1:200 dilution) along with Fc block (BD Biosciences, 1:500 dilution). GCBs (B220^+^ Fas^+^ CD38^−^) were isolated on a BD FACS Aria III sorter (BD Biosciences). Following sorting, nuclei were isolated and used for PRO-seq and PRO-cap (see below). For one experiment, ∼10^7^ GCBs from fifteen immunized mice were pooled yielding 3-4×10^6^ viable nuclei that were used for run-on. For MutPE-seq, genomic DNA was isolated from 15 x10^6^ GCBs as described previously ^117^.

### ^118^previously ^64^ with several modifications

To isolate nuclei, murine and *Drosphila* S2 cells were resuspended in cold Buffer IA (160 mM Sucrose, 10 mM Tris-Cl pH 8, 3 mM CaCl_2_, 2 mM MgAc_2_, 0.5% NP-40, 1 mM DTT added fresh), incubated on ice for 3 min and centrifuged at 700 g for 5 min. The pellet was resuspended in nuclei resuspension buffer NRB (50 mM Tris-Cl pH 8, 40% Glycerol, 5 mM MgCl_2_, 0.1 mM EDTA). For each run-on, 10^7^ nuclei of the sample and 10% *Drosophila* S2 nuclei were combined in a total of 100 µL NRB and incubated at 30°C for 3 min with 100 µL 2x NRO buffer including 5µl of each 1mM Bio-11-NTPs. In some PRO-cap experiments, the run-on reaction was performed in the presence of two biotinylated NTPs, Biotin-11-UTP and Biotin-11-CTP (Perkin-Elmer), and unlabeled ATP and GTP. Subsequent steps were performed as described (Mahat et al 2016), except that 3’ and 5’ ligations were performed at 16°C overnight and CapClip Pyrophosphatase (Biozym Scientific) was used for 5’end decapping. We also used customized adapters (PRO-seq: 3’ RNA adapter: 5’5Phos/NNNNNNNGAUCGUCGGACUGUAGAACUCUGAAC/3InvdT-3’ and 5’ RNA adapter: 5’-CCUUGGCACCCGAGAAUUCCANNNN-3’). RNA was reverse transcribed by SuperScript III RT with RP1 Illumina primer to generate cDNA libraries. Libraries were amplified with barcoding Illumina RPI-x primers and the universal forward primer RP1 using KAPA HiFi Real-Time PCR Library Amplification Kit. PRO-cap: 3’ linker (DNA oligo): 5rApp/NNNNAGATCGGAAGAGCACACGTCT/3ddC, 5’linker (RNA): ACACUCUUUCCCUACACGACGCUCUUCCGAUCUNNNNNNNNNN, reverse transcription and library amplification as in PRO-seq but using selected TruSeq_IDX_1-48 PCR primers for RT, and the same TruSeq IDX PCR primers together with TruSeq Universal Adapter forward primer for PCR. For both methods amplified libraries were subjected to gel electrophoresis on 2.5% low melting agarose gel and amplicons from 150 to 350 bp were extracted from the gel (carefully separated from possible linker dimers), multiplexed and sequenced on Illumina platforms with SR50 or longer read mode.

### RT-qPCR

RT-qPCR assays with externally spiked-in *Drosophila* S2 cells were described in detail in our previous studies ^119^. Briefly, *Drosophila* S2 cells were mixed with Ramos cells or B1-8^hi^ B cells at a ratio of 1:4 (4×10^5^ B cells and 1×10^5^ S2 cells) followed by total RNA extraction with TRIzol reagent (Thermo Fisher), DNaseI digestion, and cDNA synthesis with random primers. The 2-ΔΔCt method was used to quantify the data using the *Drosophila Act5c* transcript for normalization.

### ChIP-qPCR and ChIP-seq

ChIP-seq and ChIP-qPCR were performed as described previously without any modifications ^27,117^. Antibodies used in this study are listed in Table S1.

### ATAC-seq

ATAC-seq was performed exactly as described in detail in our previous work ^117^.

### Mutational analysis by paired-end deep sequencing (MutPE-seq)

MutPE-seq was performed following the principles described in two previous reports ^90,95^ with several adaptations. 80 ng of genomic DNA were amplified by PCR with the Kapa Hifi HS 2x RM (Roche Diagnostics), For the first PCR, we used 20-25 cycles with 0.2µM locus-specific primers fused to a varying number of random nucleotides and to the first part of the Illumina adapter sequences (FW: 5’-CTCTTTCCCTACACGACGCTCTTCCGATCT-(N)x-gene-specific sequence-3’; RV: 5’-CTGGAGTTCAGACGTGTGCTCTTCCGATCT-gene-specific sequence-3’; see Table S1 for a complete list of primer sequences).

Random Ns are used to increase complexity and shift frames of very similar amplicons to improve cluster calling and sample identification. After first PCR, the samples were purified with 0.2x/0.7x SPRI beads (Beckman Coulter), eluted in 10µl water and amplified for 10 cycles with 0.75µM primers containing linker sequences and dual barcoding (FW: 5’-AATGATACGGCGACCACCGAGATCTACAC<u>XXXXXXXX</u>ACACTCTTTCCCTACACGAC-3’; RV: 5’-CAAGCAGAAGACGGCATACGAGAT<u>XXXXXX</u>GTGACTGGAGTTCAGACGTGTG-3’ where the stretches of X dNTPs (underlined) serve as specific barcodes creating a unique dual barcode combination for each sample). PCR products were purified either by extraction from a 2% low-melt agarose gel or with 0.7x SPRI beads and eluted in 20 µl water. The concentration was determined by Picogreen-based measurements on a Nanodrop machine or by a Fragment Analyzer. Samples were equimolarly pooled for next generation sequencing on an Illumina MiSeq flowcell and sequenced paired-end (PE300).

### Primers and sgRNAs

All oligos, primers and adapters used in this study are listed in Table S1.

## Bioinformatics

### Mapping PRO-seq, PRO-cap, ChIP-seq and ATAC-seq reads to IG V regions

Reads were trimmed for standard adapters and low quality (Q<30) 3’-end bases and filtered for a remaining length of at least 20nt (excluding the UMI, if applicable) using cutadapt ^120^. Alignment to the reference genome was done with Bowtie ^121^. The NCBI GRCm38.6 assembly was used as the mouse reference. To this we added the sequence of the recombined locus containing the V region as an additional chromosome. Similarly, for human data, the Hg38 assembly was used, with the relevant V region sequence added as a chromosome. Where applicable, the dmr6 assembly of *Drosophila melanogaster* from Flybase (^122^ release 6.27) was used as the spike-in reference and was added to the alignment index. During alignment, up to three mismatches were allowed. To accommodate mapping to the repetitive *IGH/Igh* locus, a high degree of multimapping was allowed (194 potential V segments annotated for GRCm38.6). In the case of alignment with spike-ins, only reads mapping exclusively to either genome were considered. For PCR deduplication, UMIs were identified and filtered with UMI-tools (^123^ v1.0.0). No sequence differences were allowed in UMIs when collapsing the duplicates. Multimapping reads were then filtered to identify those specific to the V region sequence, taking into account the repetitiveness of the reference *IGH/Igh* locus. Reads mapping to the fixed V region sequence were allowed to multimap only to the native *IGH/Igh* locus (mouse chr12:113572929-116009954 or human chr14:105836764-106875071). The native loci were defined as the region between the earliest position belonging to an annotated V segment and the latest position belonging to an annotated J segment. Reads with a mapped location outside these areas were rejected. This filtering was implemented in a custom Python script. The qualifying reads were then classified as reads mapping uniquely to the particular V region sequence and reads mapping to both the V region sequence and to the native *IGH/Igh* region. Genome browser tracks were created by quantifying and scaling read coverage of the V region sequence using bedtools (^124^ v2-29.0).

Reads were split by strand, and strand labels for PRO-seq were inverted in order to match the strand designations of the respective PRO-cap reads. For both data types, only the first 5’ base of the strand-adjusted reads was used for the tracks. Additionally, the 5’ end of PRO-seq reads was shifted by 1 nucleotide downstream, to compensate for the fact that the last nucleotide in the RNA was incorporated during the run-on reaction. For normalized tracks, read counts were scaled to RPM (reads per million). Finally, “-“ strand coverage tracks were further scaled by –1, for visualization purposes.

### Analysis of MutPE-seq

Reads were trimmed for standard adapters with cutadapt ^120^. Poor quality (Q<25) 3’ bases were trimmed with trimmomatic ^125^ by averaging over a sliding window of 5nt. Read pairs were then filtered for minimum remaining length (200nt for read 1, 100nt for read 2) using cutadapt. Read mates were merged down to make combined single-end reads with FLASH ^126^ allowing 10% mismatch between the mates. Obvious erroneous mergers were removed by selecting combined reads with lengths within ±30nt of the amplicon length using cutadapt, The remaining combined reads were aligned with Bowtie2 ^127^, using the “–very-sensitive-local” alignment mode and only the fixed V region sequence and its immediate vicinity as reference. Samples of V genes VH4-59 and VH4-34 are different only by single nucleotide polymorphisms and were filtered for contamination by the respective second sequence by jointly aligning with the “expected” and the other “contaminating” V region sequence, discarding all aligned contaminating reads. A pile-up was generated with samtools ^128^ taking into account only bases with quality of at least 30. The pileups were then quantified with a custom Python script and the resulting mutation counts were processed and visualized with custom scripts in R (v3.5.1), with the help of additional R packages (data.table [https://cran.r-project.org/web/packages/data.table/index.html], ggplot2 [https://cran.r-project.org/web/packages/ggplot2/index.html], ggrepel [https://cran.r-project.org/web/packages/ggrepel/index.html], patchwork [https://cran.r-project.org/web/packages/patchwork/index.html]). Background mutation profiles were controlled for by subtracting the corresponding mutation frequencies in control samples from the frequencies in the samples of interest, at each position and for each substitution type. Annotation of hot and cold spots was created by means of regex search for the corresponding patterns in the reference sequence.

### Comparison on Pol II stalling and mutation frequency in V regions

Stalling sites were identified as described in Fong et al. ^91^ using the source code provided therein (https://doi.org/10.5281/zenodo.7013905) with no changes. We used the same parameters as Fong et al. to define stalling sites, namely, all sites covered by at least five PRO-seq reads and with a standard deviation three times greater than the surrounding 200 bp window ^91^. In addition, we extended these sites by 3 bp on either side to create 7 bp stalling zones. Only stalling zones within the MutPE-seq amplicon were considered and overlapping zones were merged. All other zones within the MutPE-seq amplicon were considered as non-stalling zones. This approach yielded 18-21 stalling zones and 17-19 non-stalling zones depending on the V region analyzed, which provided sufficient datapoints for statistical analyses. Mutations in all zones were counted and divided by the width of the zone to yield the normalized mutation rates which were displayed as box plots.

### Analysis of mutation frequency and nascent transcription at C:G residues

To analyze the relationship between PRO-seq read density and mutation frequency at C:G pairs (Fig. S3C-D), we first separated mutated and unmutated C:G residues into two groups. All C:G residues were extended by 3 bp on either side to create 7 bp windows. For the mutated C:G group, overlapping windows were merged. If a mutated window overlapped an unmutated one, the mutated window was retained and the unmutated window was discarded from the analysis. In this manner, only non-overlapping mutated and unmutated windows were compared. Next, the mutated windows were split into ten equal groups (deciles) ranked from lowest to highest mutation frequency. Finally, box plots were generated comparing the PRO-seq density (normalized for the length of the windows) in the unmutated and mutated groups.

In a separate analysis (Fig. S3E-F), only the 7 bp windows centered on mutated C:G residues were considered and, as above, overlapping windows were merged. Next, the PRO-seq densities in all windows were obtained. Ten equal groups (deciles) were created and were ranked from lowest to highest PRO-seq density. Box plots were generated to compare the mutation frequency of the central mutated C:G between the ten groups.

### Statistical analysis

Statistical tests were performed with the Wilcoxon rank sum test following multiple testing correction with the Bonferroni method or the unpaired Students t-test, as indicated in the figure legends.

### Data availability

All next-generation sequencing data has been deposited in GEO under accession number GSE202042. Code for the workflow and the custom scripts is available on Github at https://github.com/PavriLab/IgH_VDJ_PROcapseq and https://github.com/PavriLab/IgH_VDJ_MutPE.

### Author contributions

UES performed all PRO-seq, PRO-cap and MutPE-seq assays and analyzed data. JF generated the Ramos lines with new V regions and performed infections for MutPE-seq. KF and TN developed the bioinformatic pipeline to map NGS reads to V regions. KF developed the bioinformatic pipeline for MutPE-seq and analyzed data. RV and ITS analyzed MutPE-seq and performed the analysis of mutation and Pol II stalling in V regions. Marina M performed ChIP-seq, ChIP-qPCR and RT-qPCR in Ramos cells and analyzed data. IO performed the bioinformatic analysis of non-IG AID target genes. BB performed ChIP-qPCR and RT-qPCRs. EMW and RP generated the Eμ^−/−^ lines. MS conducted GCB PRO-seq with UES. ACG performed ATAC-seq. Marialaura M cloned JP8Bdel expression vectors for MutPE-seq. ACG performed ATAC-seq. HM provided the germline-reverted VH4-59 and VH3-30 sequences and provided feedback on the manuscript. RP conceived the project, analyzed data and wrote the manuscript with critical inputs from UES and JF.

## Acknowledgements

We thank Almudena Ramiro and Angel Alvarez-Prado (CNIC, Madrid, Spain) for sharing the mutational datasets from Ung^−/−^Msh2^−/−^ GCBs. We gratefully acknowledge the Vienna Biocenter Core Facilities (VBCF) for assistance in generating Eμ^−/−^ lines (VBCF-ProTech) and for next generation sequencing (VBCF-NGS), the IMP/IMBA BioOptics facility for flow cytometry usage, the IMP/IMBA Molecular biology services for Sanger sequencing and reagents, and the IMP/IMBA animal house.

We thank Ryan Sheridan and David Bentley (University of Colorado, USA) for advice on calling Pol II stalling sites. We thank Maximilian von der Linde for uploading NGS tracks to GEO. We thank Carrie Bernecky (ISTA, Klosterneuburg) and Clemens Plaschka (IMP, Vienna) for generating the 3D Pol II elongation complex structural visualizations. This work was funded by Boehringer Ingelheim, The Austrian Industrial Research Promotion Agency (Headquarter Grant FFG-834223), and grants from the Austrian Science Fund to UES (FWF T 795-B30) and RP (FWF P 32043-B).

## Conflicts of interest

The authors declare no conflicts of interest.

## FIGURE LEGENDS

**Figure S1:**
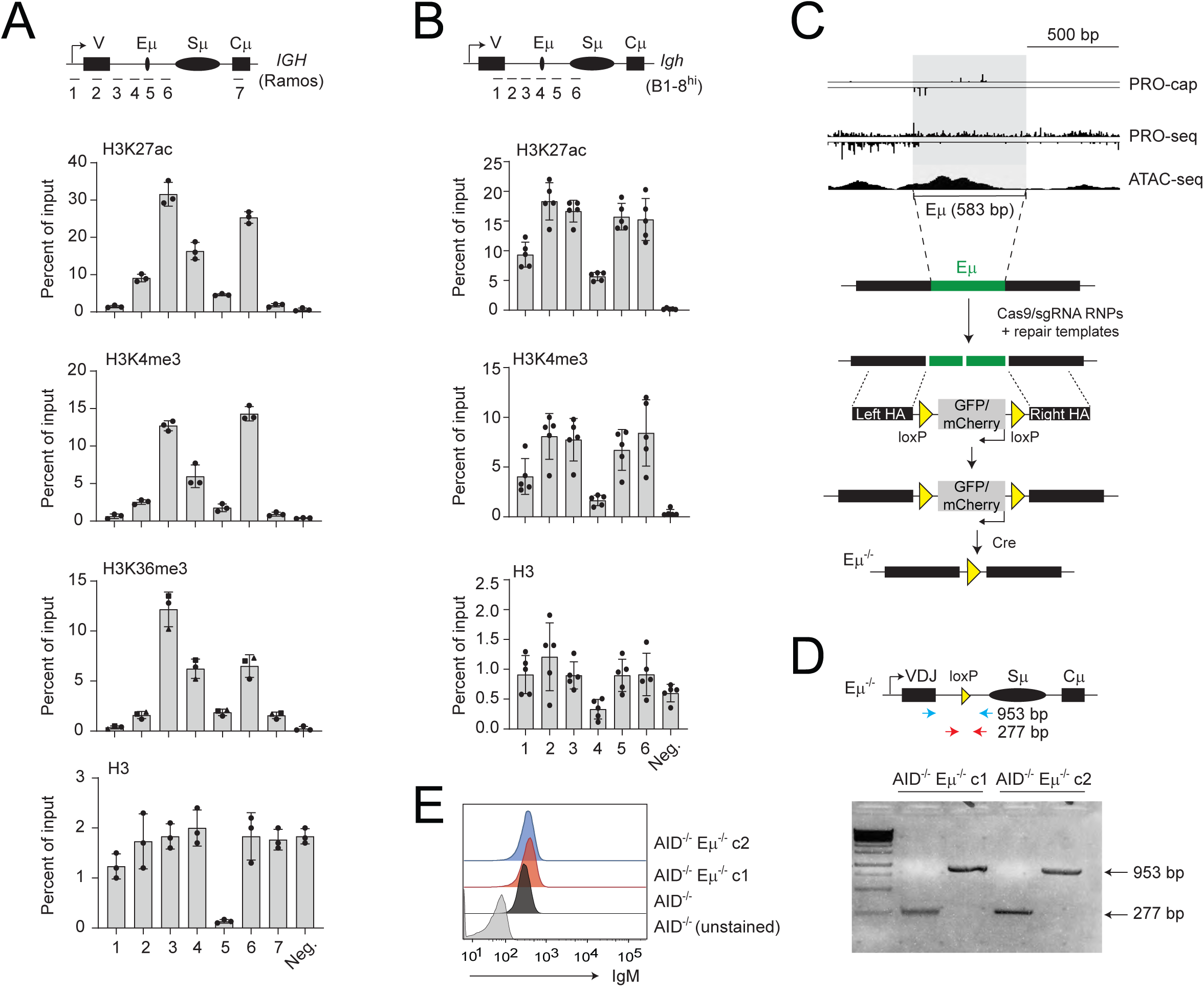
ChIP-qPCR analysis of the Ramos V region locale and generation of Ramos Eμ^−/−^ cells. (**A**) ChIP-qPCR analysis from three independent experiments in AID^−/−^ Ramos cells for the indicated epigenetic marks as well as histone H3. Amplicons (1-7) used are indicated below the locus diagram shown above the graphs. The Neg. amplicon corresponds to a gene desert on chromosome 1 and is used as a negative control. **(B)** ChIP-qPCR analysis from five independent experiments in B1-8^hi^ primary splenic cells for the indicated epigenetic marks as well as histone H3. Amplicons (1-6) used are indicated below the locus diagram shown above the graphs. The Neg. (negative region) amplicon corresponds to a gene desert on chromosome 1 and is used as a negative control. **(C)** Strategy to create the Eμ^−/−^ AID^−/−^ Ramos lines. The deleted region (583 bp) corresponds to the peak of accessible chromatin detected by ATAC-seq (top panel). This segment was replaced with a floxed reporter cassette expressing GFP or mCherry using CRISPR. Single clones double-positive for GFP and mCherry expression were isolated followed by excision of the floxed cassette by Cre recombinase. Two clones, c1 and c2 were used for all experiments. **(D)** Genotyping PCR analysis to confirm the loss of Eμ in AID^−/−^ Eμ^−/−^ clones c1 and c2. The location of primers is shown in the diagram above the gel image. **(E)** Surface IgM expression in Eμ^−/−^ AID^−/−^ Ramos clones relative to the parental AID^−/−^ line determined by FACS.

**Figure S2:**
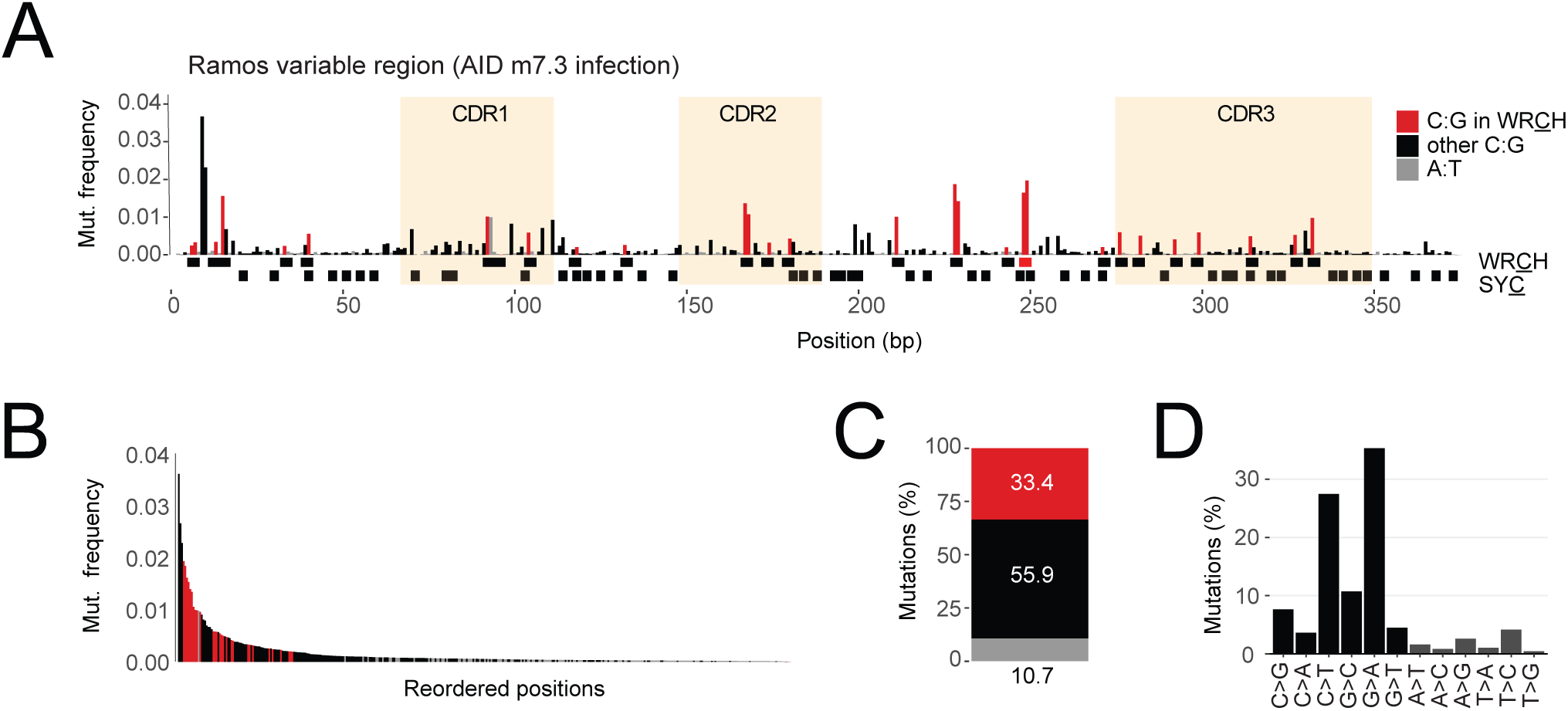
MutPE-seq of the Ramos V region following infection of AID^−/−^ cells with AID (m7.3). (**A**) Mutation profile of the Ramos V region generated by AID (m7.3) following 21 days of infection. The type of mutation and location of hotspots is indicated and can be compared directly with Fig. 3E. **(B—D)** Waterfall plot of mutations ordered by frequency (B), bar plot of mutation frequencies in the three indicated classes (C) and bar graph of mutation type (D), as explained in the legend of Fig. 3 F to H).

**Figure S3:**
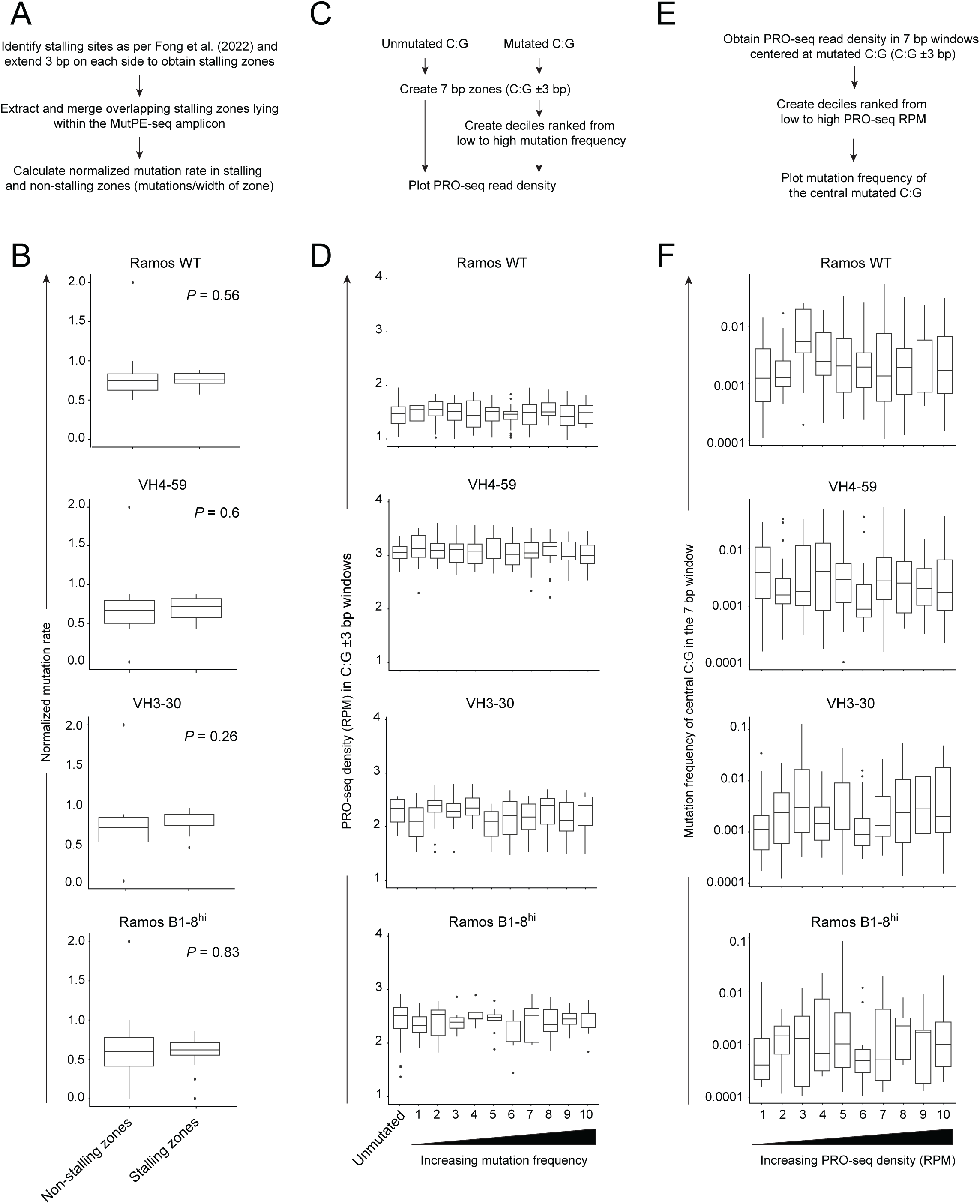
Relationship between SHM, Pol II stalling and nascent transcription at different V regions. **(A)** Scheme for the analysis of mutation frequency between stalling sites and stalling zones within V regions. **(B)** Box plots comparing mutation frequency between stalling and non-stalling zones in the Ramos endogenous (WT) V region, VH4-59, VH3-30 and Ramos^B1–8hi^. Statistical analyses were performed with a Wilcoxon rank sum test and indicated within the plots. **(C)** Categorization of unmutated and mutated C:G pairs from MutPE-seq data, as described in the text. **(D)** Box plot showing the PRO-seq density in 7 bp windows centered at each C:G residue in the unmutated group and the ten mutated sub-groups, as described in A. The V gene analyzed is indicated at the top of each plot. The medians are indicated by black lines inside the boxes. A Wilcoxon rank sum test after multiple testing correction using the Bonferroni revealed no statistical significance (defined as *P* < 0.05) between any pair of groups within any of the box plots. **(E)** Workflow for the classification of mutated C:G neighborhoods into 7 bp windows (C:G ±3 bp) based on their PRO-seq levels, as described in the text. **(F)** Box plots showing the mutation frequencies of the central C:G residue in the 7 bp windows (C:G ±3 bp) within all ten PRO-seq sub-groups. The V gene analyzed is indicated at the top of each plot. The medians are indicated by black lines inside the boxes. A Wilcoxon rank sum test with the Bonferroni correction revealed no statistical significance (defined as *P* < 0.05) between any pair of groups within any of the box plots.

**Figure S4:**
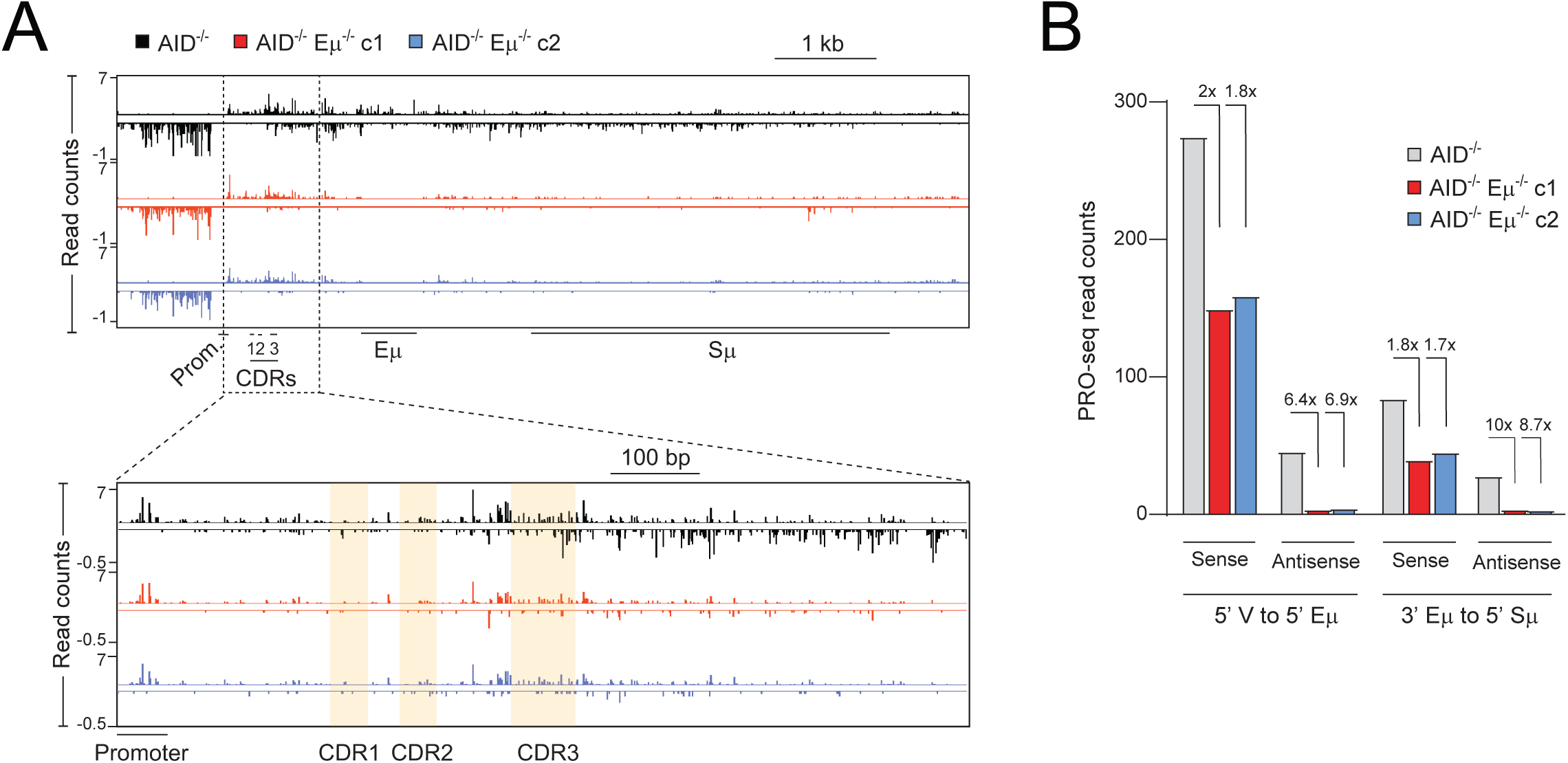
PRO-seq profiling of the Ramos V region in AID^−/−^ Eμ^−/−^ cells. **(A)** Analysis of nascent transcription 3’ ends (PRO-seq) at the *IGH* locus in AID^−/−^ and AID^−/−^ Eμ^−/−^ Ramos cells (c1, c2). The promoter (Prom.), complementary determining regions (CDR 1-3), intronic enhancer (Eμ) and switch µ region (Sμ). The lower panel shows a magnified view of the V region locale. **(B)** Quantification of PRO-seq read counts from the data in A. Signals were quantified separately on the sense and antisense strands and further divided into regions upstream and downstream of the deleted Eμ segment. The upstream portion extends from the 5’ end of the V region to the 5’ end of the deleted Eμ segment. The downstream portion extends from the 3’ end of the deleted Eμ segment to the 5’ end of Sμ.

**Figure S5:**
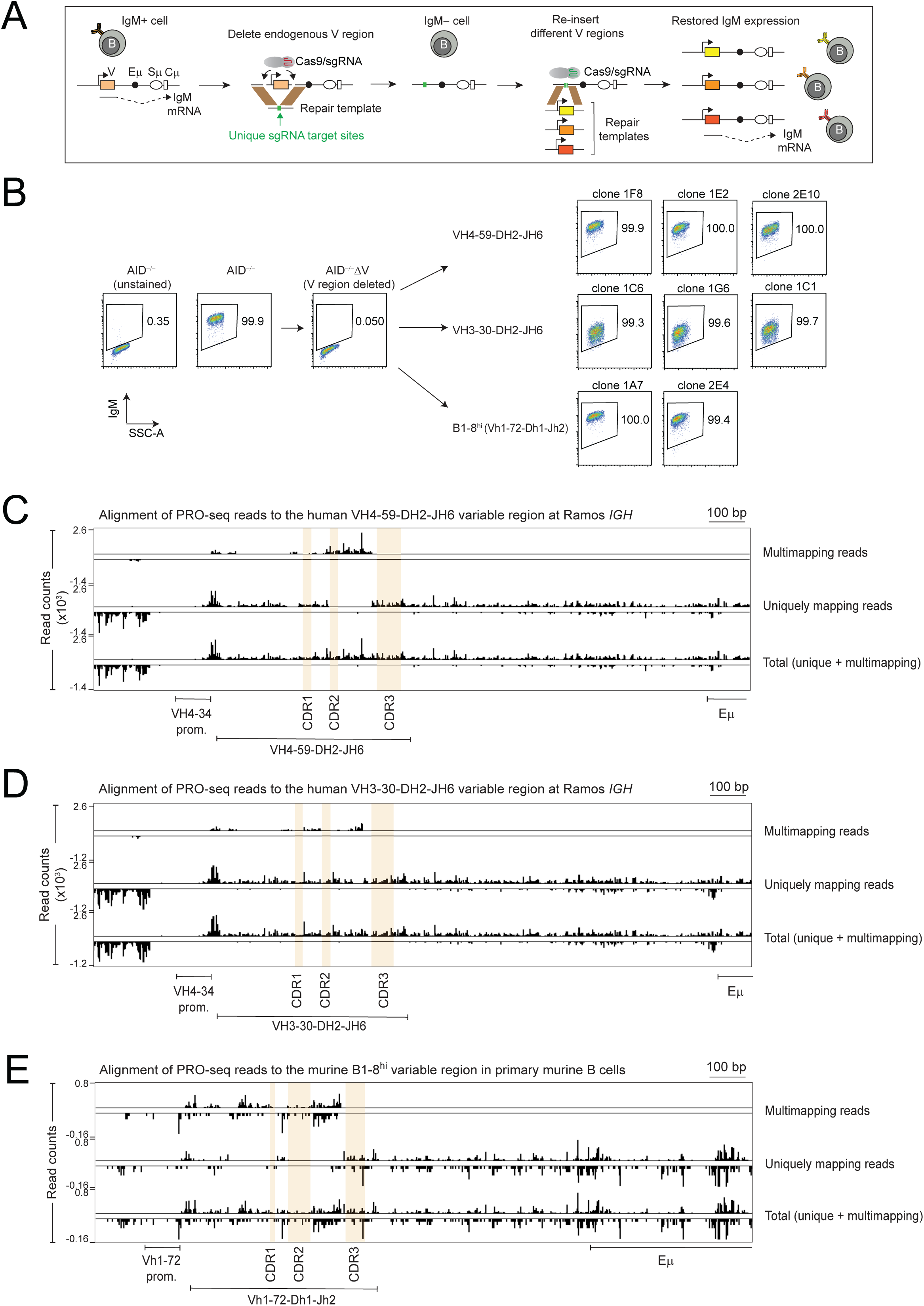
Generation of Ramos cell lines expressing exogenous V regions. **(A)** Schematic representation of the workflow for generating Ramos cells expressing new V regions using CRISPR-based editing. An IgM-negative line was made by excising the endogenous V region in AID^−/−^ cells (AID^−/−^ΔV) and replacing it with a unique small guide RNA (sgRNA)-targeting sequence (green). Subsequently, an sgRNA targeting this site is combined with Cas9 and homology repair templates harboring any new V regions. Correct integration leads to restoration of surface IgM expression which is used as a readout to identify correctly targeted clones. **(B)** Flow cytometry analysis of surface IgM expression in Ramos cell clones expressing the indicated human and mouse V regions. Shown are three clones of human VH4-59-DH2-JH6 and human VH3-30-DH2-JH6 expressing Ramos cells, and two clones of B1-8^hi^ expressing Ramos cells. **(C)** PRO-seq analysis showing the 3’ ends of aligned reads at the human VH4-59-DH2-JH6 V region expressed from the VH4-34 promoter at the human *IGH* in Ramos cells. Tracks of Multimapping, uniquely mapping and total are shown (see also Fig. 3A and 3B for a detailed description). **(D)** PRO-seq analysis as in C of the human VH3-30-DH2-JH6 V region expressed from the VH4-34 promoter at the human *IGH* in Ramos cells. **(E)** PRO-seq analysis as in C at the murine B1-8^hi^ V region at the *Igh* locus in mice.

**Figure S6:**
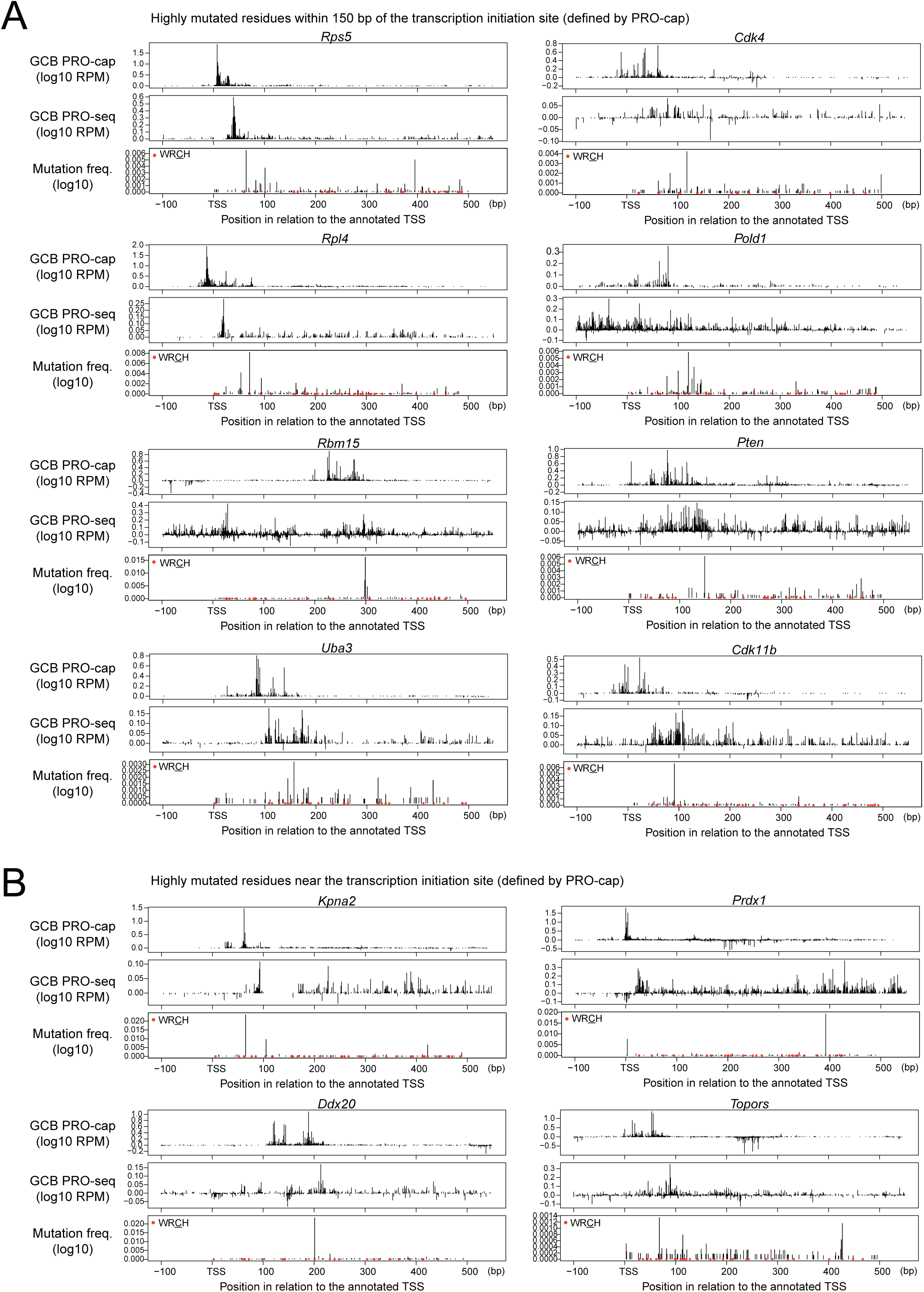
Analysis of PRO-cap, PRO-seq and SHM at non-Ig AID target genes in murine GCBs. **(A)** 5’ and 3’ ends of nascent RNA (PRO-cap, PRO-seq) and mutation profiles (from Alvarez-Prado et al. ^17^) at selected AID target genes as in Fig. 5D-G. Shown are genes where highly mutated residues lie within 150 bp of the transcription initiation site defined by the peak of PRO-cap signals. The WR<u>C</u>H motifs are indicated as red dots. (B) As in A but showing genes where highly mutated residues lie near the transcription initiation site defined by the peak of PRO-cap signals.

**Figure S7:**
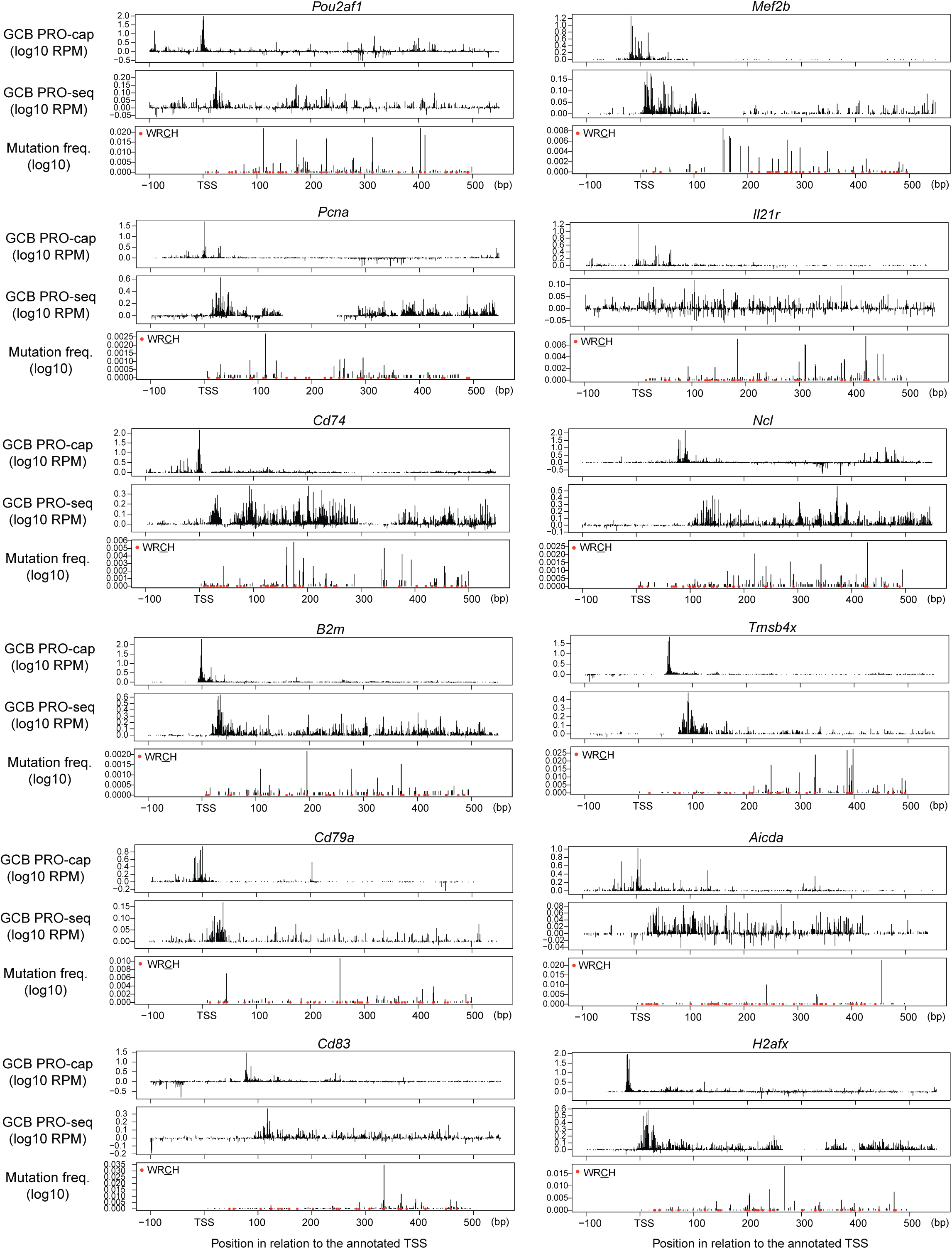
Additional examples of PRO-cap, PRO-seq and SHM at non-Ig AID target genes in murine GCBs. See Fig. 5D for detailed legend.

**Figure S8:**
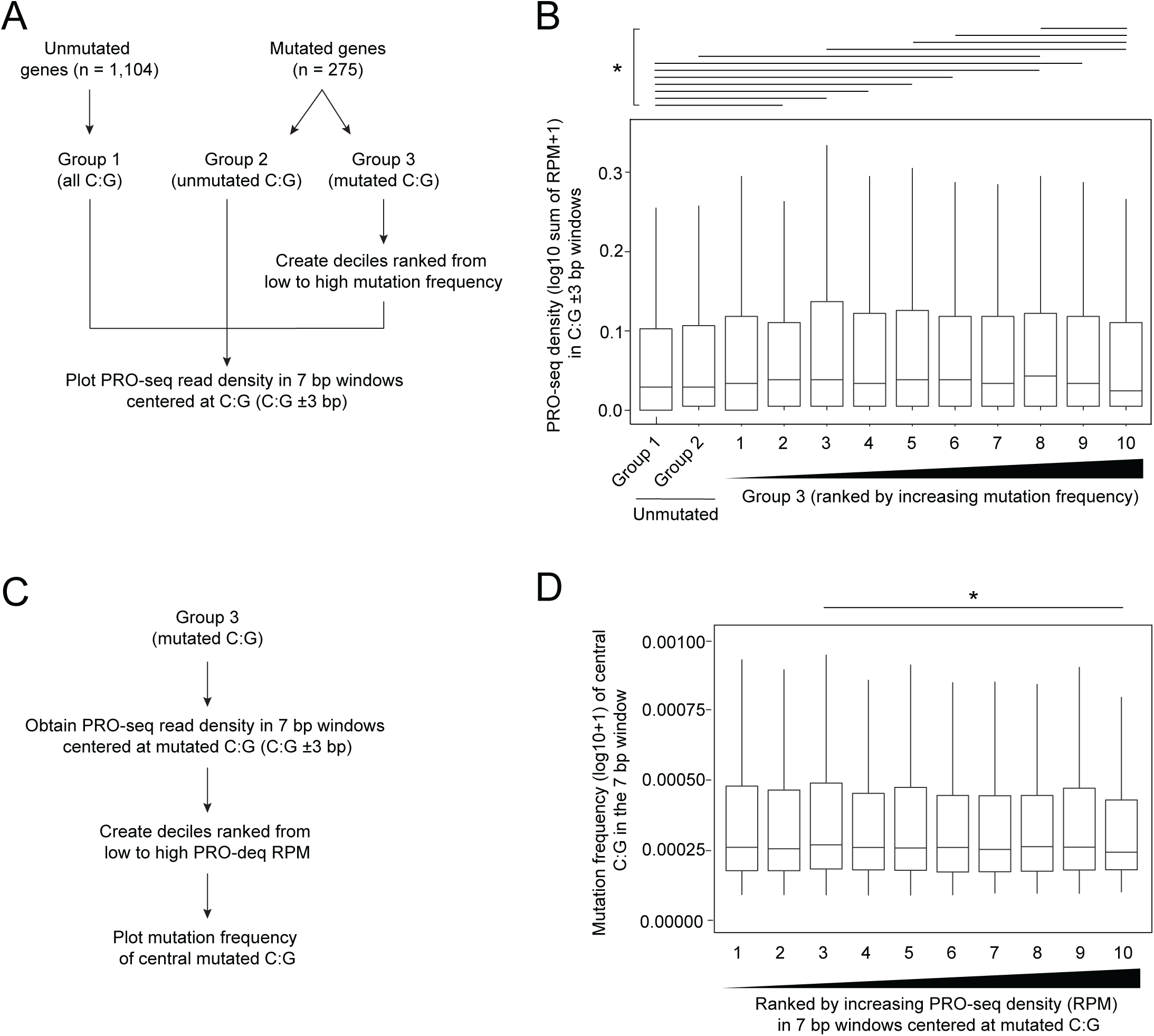
Relationship between SHM and transcriptional strength at non-IG AID target genes. (**A**) Workflow for the classification of mutated and unmutated genes. **(B)** Box plot showing the PRO-seq density in 7 bp windows centered at each C:G residue in the three groups and sub-groups described in A. The medians are indicated by black lines inside the boxes. The asterisk indicates *P* < 0.05 based on the Wilcoxon rank sum test after multiple testing correction using the Bonferroni method. **(C)** Scheme for the classification of mutated C:G neighborhoods based on their PRO-seq density. **(D)** Box plots showing the mutation frequencies of the central C:G residue in the 7 bp windows (C:G ±3 bp) within all ten PRO-seq sub-groups. The medians are indicated by black lines inside the boxes. The asterisk indicates *P* < 0.05 based on the Wilcoxon rank sum test after multiple testing correction using the Bonferroni method.

**Table S1:**
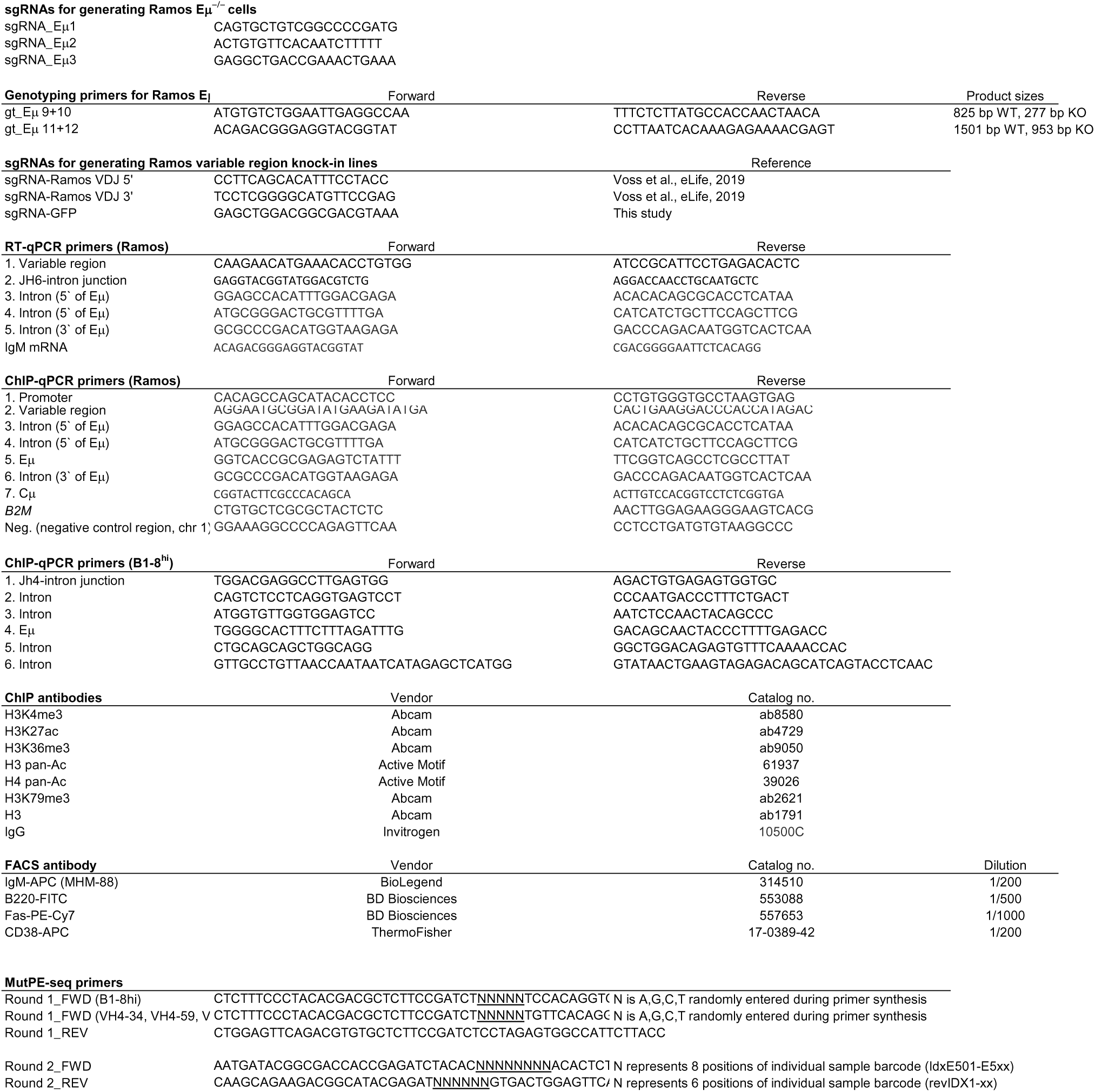
List of all primers, oligos, antibodies and sgRNAs used in this study.

**Table S2:**
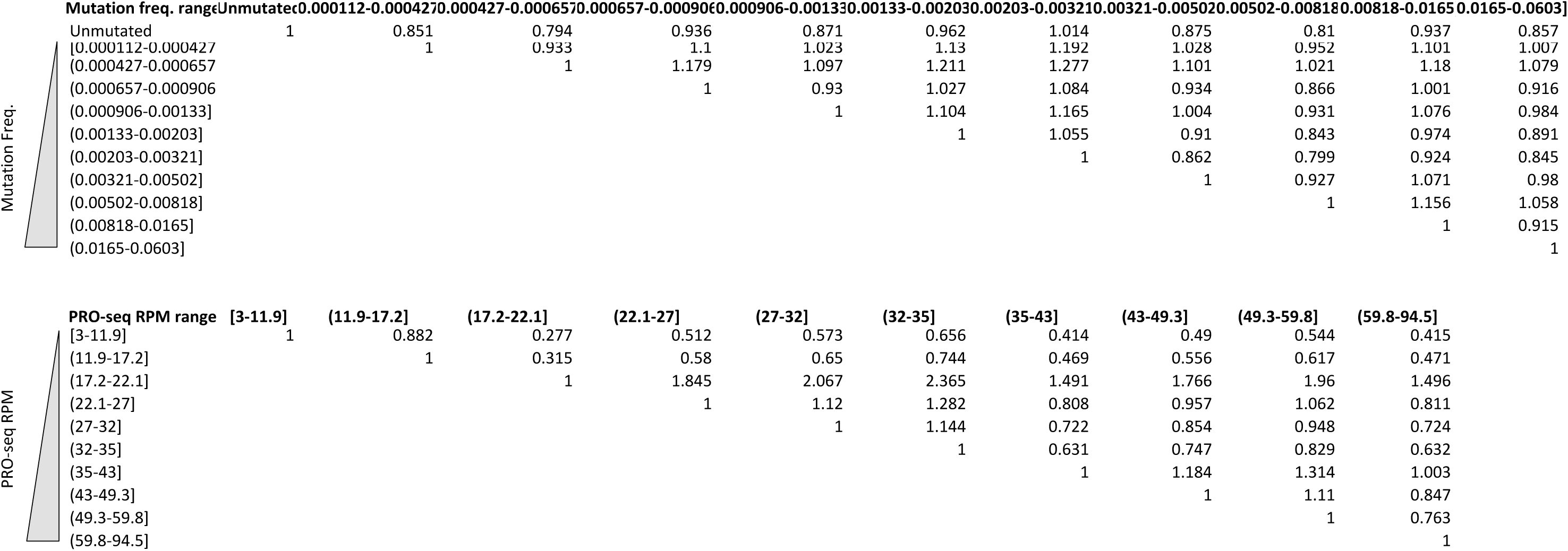
Statistical analysis of mutation frequencies and PRO-seq densities in different V regions. he upper and lower tables show the *P* values for the analyses in Fig. S3D and S3F, respectively. *P* values are based on the Wilcoxon rank sum test following multiple testing correction with the Bonferroni method.

## Notes

### Competing Interest Statement

The authors have declared no competing interest.

### Summary of Updates

The text has been revised in several sections for clarity. Figure S7-8 and Tables have been revised. Author affiliations updated.

